# Human Organoid Tumor Transplantation Identifies Functional Glioblastoma - Microenvironmental Communication Mediated by PTPRZ1

**DOI:** 10.1101/2024.05.02.592055

**Authors:** Weihong Ge, Ryan L. Kan, Can Yilgor, Elisa Fazzari, Patricia R. Nano, Daria J. Azizad, Matthew Li, Joyce Y. Ito, Christopher Tse, Hong A. Tum, Jessica Scholes, Kunal S. Patel, David A. Nathanson, Aparna Bhaduri

## Abstract

Glioblastoma, the most aggressive and deadly form of primary brain cancer, is driven by both intrinsic cellular properties and external factors from the tumor microenvironment. Here, we leverage our novel human organoid tumor transplantation (HOTT) system to explore how extrinsic cues modulate glioblastoma cell type specification, heterogeneity, and migration. We show that HOTT recapitulates the core features of major patient tumor cell types and key aspects of peritumor cell types, while providing a human microenvironment that uniquely enables perturbations in both the patient tumor and its microenvironment. Our exploration of patient tumor – microenvironmental interactions in HOTT highlighted PTPRZ1, a receptor tyrosine phosphatase implicated in tumor migration, as a key player in intercellular communication. We observed that tumor knockdown of PTPRZ1 recapitulated previously described roles in migration and maintaining progenitor identity. Unexpectedly, environmental PTPRZ1 knockdown drove opposite migration and cell fate changes in the tumor, even when the tumor was not manipulated. This previously undiscovered mode of tumor-microenvironmental communication highlights the need to study human glioblastoma in the context of a human microenvironment such as HOTT.

## Introduction

Glioblastoma (GBM) is an aggressive form of adult brain cancer, with limited treatment options and average survival of 12 - 18 months post diagnosis (Omuro and DeAngelis, 2013; Wen et al., 2020). Molecular profiling of GBM has long shown inter- and intra-tumoral cell type heterogeneity (Neftel et al., 2019b; Ravi et al., 2022; Tirosh et al., 2016; Venteicher et al., 2017), with this diversity of cell types possibly contributing to difficulty in developing effective treatments (Couturier et al., 2020; Neftel et al., 2019b). Cell types identified in GBM include ones reactivated from cortical development such as outer radial glia (Bhaduri et al., 2020b; Wang et al., 2020). Outer radial glia are a subtype of neural stem cells (Hansen et al., 2010; Pollen et al., 2015) abundant in the developing human cortex, in contrast to the rodent where they are exceptionally sparse (Lui et al., 2011). Previous research also suggests that other subtypes in the tumor may also reflect human neurodevelopmental programs (Bhaduri et al., 2020b; Couturier et al., 2020), emphasizing the emerging need to study GBM in human patient relevant model systems.

Numerous models are commonly used to study GBM. These include immortalized cell lines, low passage patient derived gliomaspheres, orthotopic xenografts, and organoid models. Some of these cell culture based models are enriched for glioma stem-like populations (Lathia et al., 2015), and thus do not fully recapitulate the tumor heterogeneity (Pastrana et al., 2011). Novel approaches with organoid technologies have expanded the opportunity to interrogate the three-dimensional tumor context and facilitate direct leveraging of patient samples. For example, recent innovations in explant technologies include the glioblastoma organoid (GBO) (Jacob et al., 2020) and patient derived explant (PDE) systems (LeBlanc et al., 2022). These approaches allow for the *in vitro* examination of patient tumors and maintain much of the patient tumor heterogeneity. Other patient derived approaches such as the cerebral organoid glioma (GLICO) (Linkous et al., 2019) model require initial plating of glioma stem cells from the patient tumor. In our previous work (Bhaduri et al., 2020b), we developed a direct from patient model that enables tumor cell dissociation from freshly resected tumors followed by immediate transplantation into a human cortical organoid model. These models are excitingly opening the frontier of studying human tumors in systems that do not require immortalization and thus compromising of potentially relevant heterogeneous patient tumor cell characteristics.

As models to study GBM have expanded, so has our knowledge of how they interact with their microenvironment. Recent studies have shown that GBM cells interact with normal cell types, such as neurons and astrocytes, through a variety of mechanisms. For example, brain tumors are increasingly being observed to electrically and synaptically integrate into brain circuits (Krishna et al., 2023; Venkatesh et al., 2019), leveraging properties of neurons to promote invasion through the brain (Venkataramani et al., 2022) and being interconnected into functional networks through electrical activity and microtube connections (Osswald et al., 2015). Together, the study of the tumor microenvironment (TME) has evolved into the new field of cancer neuroscience (Mancusi and Monje, 2023; Winkler et al., 2023), highlighting that unique features of the brain make this TME specifically conducive to GBM biology. However, given that the human cortex is markedly different than the rodent in terms of cell types and structural features (Bakken et al., 2021), many open questions exist regarding how the interactions between the human brain and heterogeneous GBM cell types promote key aspects of brain tumor biology. These questions include understanding how tumor - TME interactions drive tumor growth, but also how these interactions mediate cell fate specification and tumor migration.

It is well understood that during human cortical development, cell fate specification is a crucial process in generating the diverse cell types that ultimately comprise cortical layers and distinct functional areas (Cadwell et al., 2019). Given that a subset of these cell types are reactivated in GBM, understanding cell fate regulation in this context is additionally essential to understanding tumor progression. Indeed, tumor cell types change over the progression of the disease or across contexts (Neftel et al., 2019b), sometimes in response to treatments such as radiation (Gupta and Burns, 2018; Muthukrishnan et al., 2022; Pajonk et al., 2010) and in other cases as recurrent tumors that emerge with more aggressive cell types (Wang et al., 2022). Normal development is governed by a tightly regulated set of intrinsic and extrinsic factors that influence this cell fate specification and also behavior of dividing and migrating cells; we reasoned that it would be important to explore these features in the context of GBM and a human TME.

Our overarching aim is to establish and interrogate a model of how the tumor and the TME reciprocally interact to promote tumor migration and cell fate specification. Existing organoid transplantation models have relied on established organoid growth conditions for tumor growth (Bhaduri et al., 2020b; Linkous et al., 2019), but have not been optimized in the same way that two-dimensional cell line culture has been developed (Hasselbach et al., 2014). Recognizing that the status quo might not be the most ideal condition for replicating the heterogeneity and molecular features of patient tumors, we decided to explore microenvironmental conditions in our human organoid tumor transplantation model (HOTT) beginning with the characterization of different media conditions to establish optimal conditions. We found that not only does HOTT recapitulate patient tumor characteristics, but it also recapitulates important features of the peritumor environment cell types that differ from the healthy adult brain. Thus, HOTT is an ideal model to understand cell - cell relationships between the tumor and its microenvironment. These efforts gave us confidence in how the HOTT system recapitulates tumor heterogeneity and cell type fidelity, identifying novel tumor - TME interactions mediated by an outer radial glia marker gene, PTPRZ1, in driving tumor migration and cell fate specification changes. Although we validated, as previously described, that PTPRZ1 is required for tumor migration and maintenance of stemness within GBM cells, we also discovered that environmental knockdown of PTPRZ1, without modifying the tumor directly, results in increased tumor migration and stemness. This is a previously undescribed mechanism by which the tumor and the TME communicate and a novel description of PTPRZ1 biology that is independent of its catalytic activity. Together, this study emphasizes the value of the HOTT system as a tool to study and manipulate TME interactions, and further emphasizes the need to contextually study human GBM biology within a TME.

## Results

### Human Organoid Tumor Transplantation (HOTT) is a valuable tool to study intrinsic and extrinsic regulators of GBM biology

We use our HOTT system to explore both intrinsic and extrinsic features of five GBM patient tumors (Fig 1A, STable 1). This approach closely mimicked the direct from patient organoid transplantation system we have previously described (Bhaduri et al., 2020b). Each of these tumors was dissociated and labeled with a lentivirus expressing green fluorescent protein (GFP), and the input tumor was analyzed with single-cell or single-nucleus sequencing. As the substrate for our GBM cell transplantation, we used human embryonic stem cell (hESC) derived cortical organoids generated using established protocols (Bhaduri et al., 2020a; Kadoshima et al., 2013; Pollen et al., 2019). To maximally leverage the cell type heterogeneity present in the cortex, while ensuring the consistency of our organoid pipeline for tumor transplantations, we chose to perform the transplantations at stages after peak neurogenesis (SFig 1A). Although organoids are a developmental model, we and others have shown that they robustly support patient GBM cell type heterogeneity (Bhaduri et al., 2020b; Linkous et al., 2019) and here we observed that they also mimic important aspects of the TME as well, making them an ideal tool to study of tumor - TME interactions.

**Fig 1:**
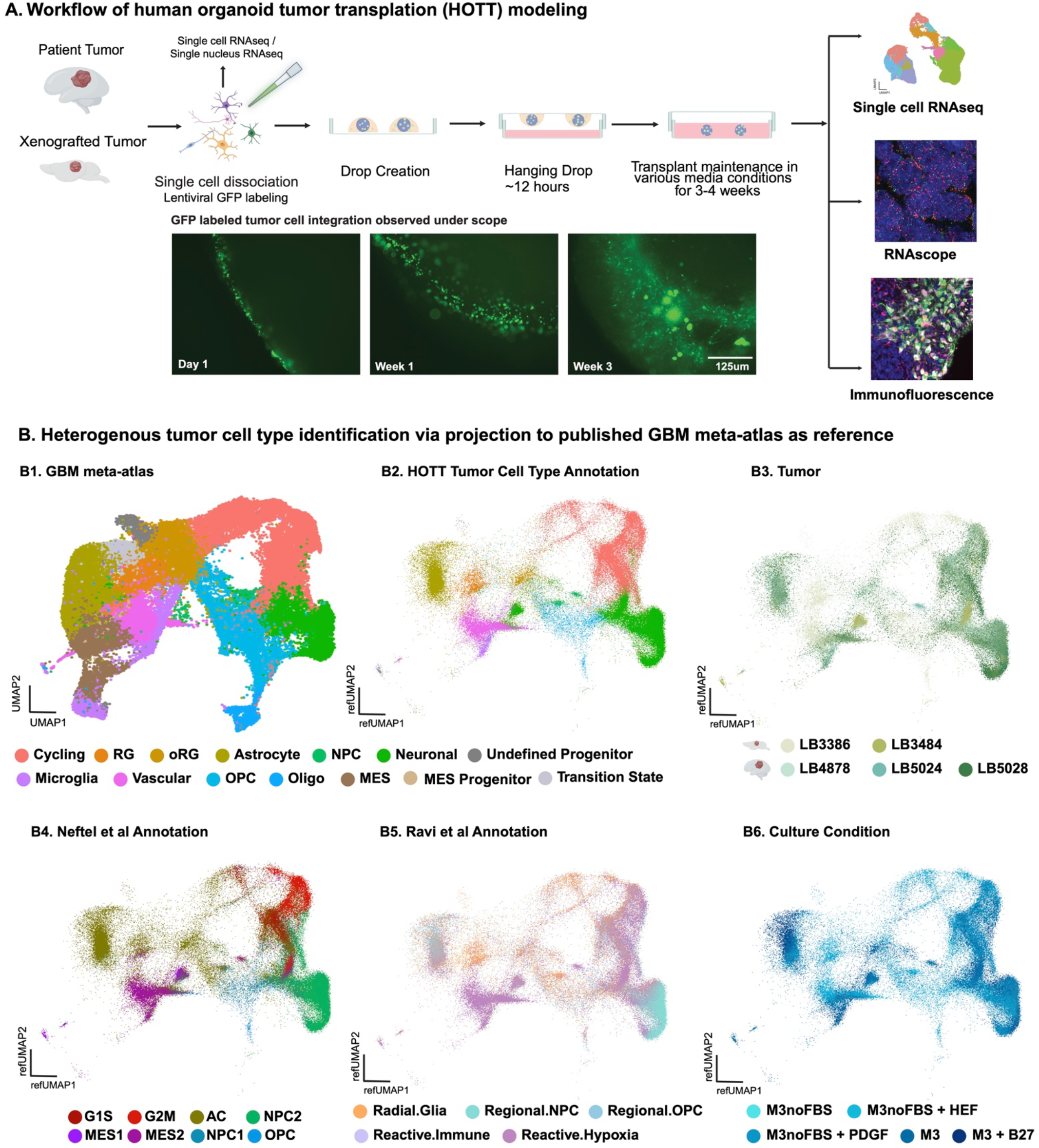
Establishment of Human Organoid Tumor Transplantation (HOTT) model. **A)** Workflow of <u>H</u>uman <u>O</u>rganoid <u>T</u>umor <u>T</u>ransplantation (HOTT) modeling. Tumors were obtained from either freshly resected surgical sampling or from dissection of patient xenografted mice. These tumors were then dissociated, and part of the original tumor was sequenced with single-cell or single-nuclei transcriptomics. The rest of the tumor was transplanted onto cortical organoids via a novel hanging drop method after GFP labeling. The organoids were then maintained in different media conditions to explore various environmental conditions for 3-4 weeks before harvest. Half of the organoids were processed via fluorescent activated cell sorting (FACS) to isolate and sequence GFP+ tumor cells, GFP-organoid cells, and tumor-naïve batch-matched organoids, while the other half of transplanted organoids were used for immunofluorescence and RNAscope validation. The experimental workflow was implemented on three primary tumors (indicated by the human brain logo) and 2 direct-from-patient orthotopic xenograft (DPDOX) tumors (represented by the mouse brain logo). **B)** The entire set of experimental media conditions across five tumor samples was analyzed jointly. Cell type annotation was performed by projection to a compendium of GBM published dataset **(B1),** these cell types were well represented and heterogeneous GBM cell types were identified **(B2),** across 5 tumors **(B3)** with a robust correlation with annotations from Neftel et al **(B4)** and annotations from Ravi et al **(B5).** No dramatic separation 5 culture conditions **(B6)** in UMAP space was detected.

To optimize tumor engraftment, we tested multiple methods of tumor transplantation and observed that our unique hanging drop method resulted in the highest engraftment efficiency (SFig.1B, see Methods). Using this engraftment strategy, we subjected each tumor implanted in the HOTT system to 5 different media conditions (SFig.1C). These media conditions were designed based upon commonly utilized gliomasphere growth methods and with the knowledge that certain media components such as FBS have been historically associated with poor growth and differentiation (Lee et al., 2006). As a baseline, we used our standard organoid ‘Media 3’ (M3) (Methods). We included a second condition using our standard organoid media minus FBS, predicting this would drive more progenitor populations. Because gliomasphere cultures are often supplemented with growth factors (Lee et al., 2006), a third condition supplemented fibroblast growth factor (FGF) and epidermal growth factor (EGF) to this FBS depleted media. To test the hypothesis that distinct growth conditions would yield unique fractions of progenitor populations in the tumor, we also included a condition supplementing platelet derived growth factor (PDGF), a ligand for PDGFRA-expressing oligodendrocyte precursor cells (Zhu et al., 2014). Finally, we include a condition where we added B27 to test whether this may generate more tumor derived neuronal populations (SFig 1C).

After 3-4 weeks in culture, we isolated the tumor cells using fluorescent activated cell sorting (FACS) (SFig 1D) and performed single-cell RNA-sequencing on the GFP positive tumor cells as well as a subset of the GFP negative normal organoid cells and paired tumor naive organoids from the same differentiation batch yielding a total of 216,125 single cells passing quality control (STables 2 – 3). In parallel, HOTTs were fixed for immunofluorescence and fluorescent in situ hybridization assays (FISH, using RNAScope). The sequenced cells were analyzed for copy number variation (CNV) (Tickle T, 2019) (SFig 1E), accounting for any false positive or negatively sorted cells. Initial analyses were performed with the tumor cells only, and annotations were performed by projecting our collected data onto a reference meta-atlas that is a compendium of published datasets (Bhaduri et al., 2020b; Couturier et al., 2020; Darmanis et al., 2017; Jacob et al., 2020; Neftel et al., 2019a; Yu et al., 2020; Yuan et al., 2018) benchmarked against existing annotations commonly used in the literature (Fig 1B, SFig 2A). Across tumors, we observed a strong representation of most GBM cell types, with expected limited representation of mature oligodendrocyte, mesenchymal and immune related programs (Fig 1B, SFig 2B). When examining media conditions and tumor of origin (Fig 1B), we observed that the 5 tumors overlapped in UMAP space. These initial analyses suggest that HOTT recapitulates tumor heterogeneity and cell types that have been previously described in human GBM.

### HOTT cell type composition is preserved across environmental conditions

Given there was excellent representation of most cell types across tumors, we wanted to quantify the cell type composition for each tumor and media condition (Fig 2A, SFig 2C). Overall, we found that most media conditions had relatively consistent cell type distributions, with the exception that there were more progenitors in the FGF and EGF supplemented condition, which is consistent with prior literature (Lee et al., 2006) and our own expectations. By both single-cell analysis and immunofluorescence for stemness markers NES and SOX2 (Fig 2B), we observed the preservation of tumor progenitor cells, such as neural progenitor cells and outer radial glia, across media conditions. Notably, these cell types were present even in culture conditions that contained FBS, which is notorious for inducing neural stem cell differentiation and thereby losing progenitor populations that could give rise to cancer stem cells (Janiszewska et al., 2012; Wakimoto et al., 2009).

**Fig 2.**
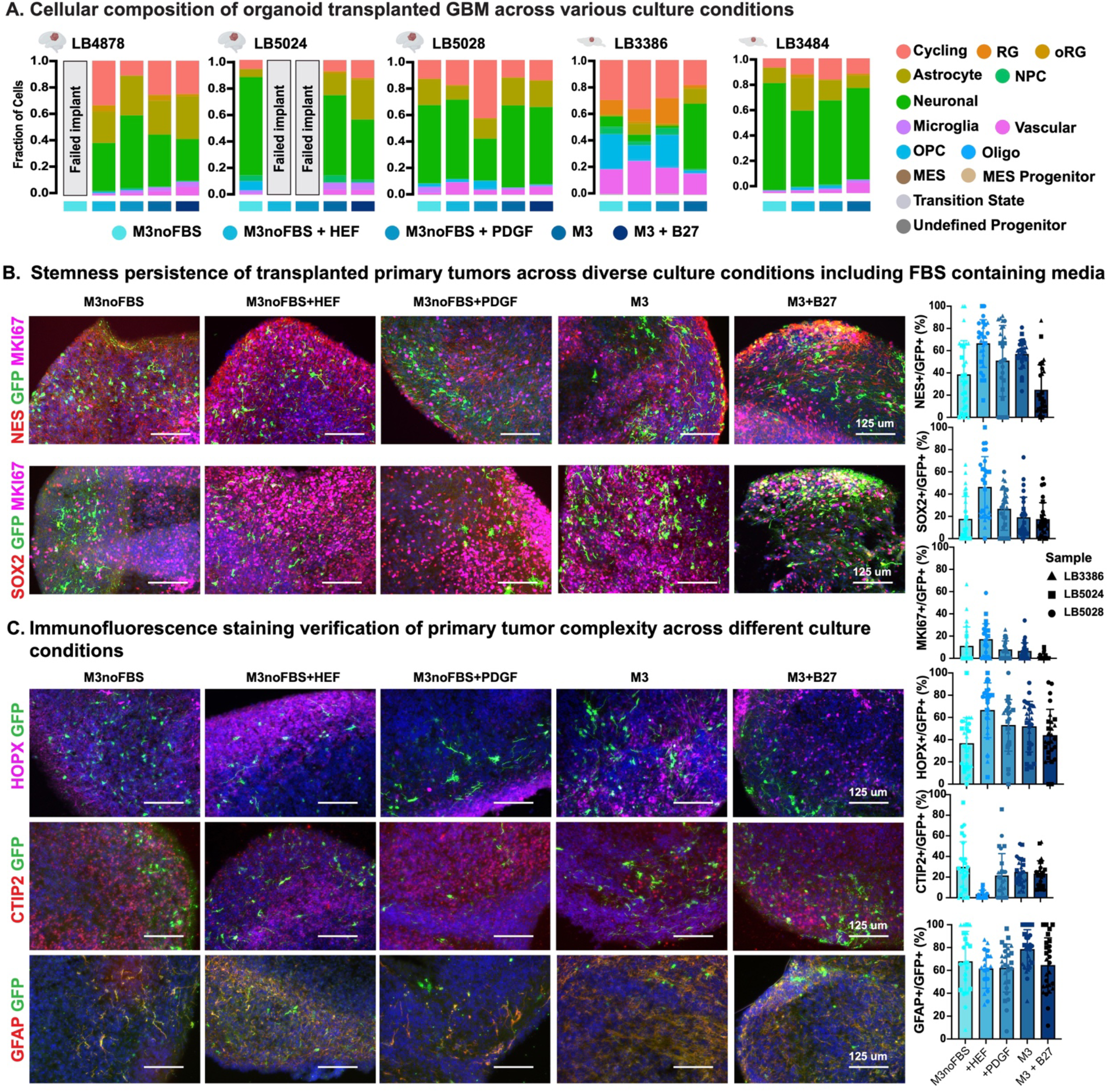
Transplanted tumors maintain stemness and preserve primary tumor complexity. **A)** Cellular composition of organoid transplanted GBM across various culture conditions. HOTT modeling enables robust engraftment of patient tumors. Notably, 100% success engraftment is achieved in FBS containing culture conditions. In addition, the intricate complexity of primary tumors is also preserved across all media conditions. Stacked bar charts are shown representing the fraction of each cell type identified via single-cell analysis for each tumor and condition. **B)** Transplanted primary tumors across diverse culture conditions demonstrated persistence of their stemness features as measured by MKI67+, SOX2+ and NES+ immunostaining and their quantification as a fraction of the GFP+ population across multiple tumors. This stemness was observed across all media conditions, including those containing FBS. **C)** Verification of primary tumor complexity across various culture conditions was performed using cell type-specific marker staining, including the progenitor marker HOPX, neuronal markers CTIP2, as well as the astrocyte marker GFAP. The accompanying bar graphs, representing multiple tumor samples, illustrate the prevalence of the most populated cell types across different culture conditions. Notably, the culture condition without FBS supplemented with EGF and bFGF (i.e., M3noFBS+HEF), preferred for gliomasphere growth, demonstrated a higher abundance of dividing cells and progenitors, while exhibiting fewer neuronal cells. This observation is further supported by single-cell RNAseq data, which indicated a higher proportion of cycling and radial glial (RG) populations **(2A)**

We validated the cell type distributions observed via single-cell RNA-sequencing with immunofluorescence analysis for cell type marker genes. Notably, across media conditions, most markers including HOPX (outer radial glia), CTIP2 (deep layer neurons), and GFAP (astrocytes) were consistent across media conditions, though we also observed a significantly increased progenitor compartment in FGF and EGF media conditions (Fig 2C, SFig 2D). However, across conditions, the only media condition in which engraftment was achieved across all replicates was in media conditions including FBS in the media (Fig 2A, SFig 2C). Across media conditions, the number of GFP positive cells identified via immunofluorescence matched the number of single cells retrieved for the analysis (SFig 2E), with the highest numbers in standard organoid media and the FBS depleted, FGF and EGF supplemented condition. These data suggested that progenitor cells are maintained across media conditions, and that although FBS depleted, FGF and EGF supplemented media promoted the most progenitor cells, standard organoid growth conditions were, in fact, the most reliable.

### HOTT shows fidelity to primary tumor cell types and the cortical TME

Given that tumor heterogeneity could be observed across media conditions in the HOTT system, we sought to explore whether the heterogeneity and cell type fidelity was altered in the HOTT system compared to the primary matched tumors as well as our meta-atlas of published primary molecular profiles. We first looked for how the cell types across media conditions compared to patient tumor heterogeneity by leveraging a correlative analysis in which we compared clusters from individual HOTT media conditions to their matched patient cell types (Fig 3A). We observed that not only were diverse patient cell types represented across media conditions, but also many clusters were highly correlated to the original patient tumor, indicating tumor heterogeneity is well preserved in HOTT. Similar observations were made when comparing to other published annotations (SFig 3A-B), suggesting that regardless of the granularity of the annotation, HOTT recapitulates patient tumor heterogeneity.

**Fig 3.**
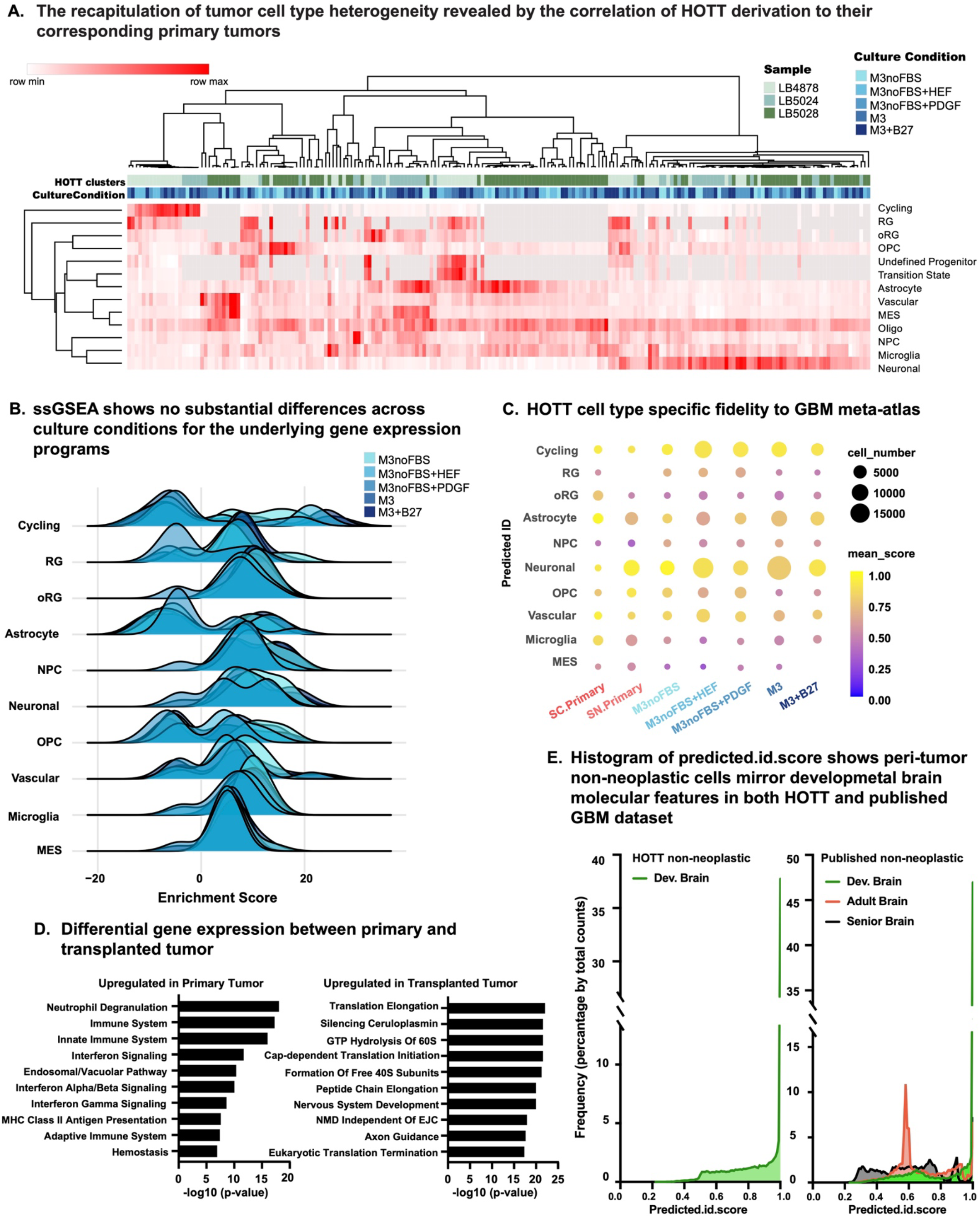
HOTT derived tumor cells recapitulate the molecular features of their corresponding primary tumors. **A)** Three HOTT samples were individually clustered using the Seurat pipeline, and resulting clusters were merged and correlated to their corresponding primary tumor cell types annotated by projection onto the GBM meta-atlas. The correlation to their corresponding primary tumors demonstrates the recapitulation of tumor cell type heterogeneity. **B)** ssGSEA analysis was executed to determine whether fidelity to individual cell types was altered across media conditions. Ridge plots of each media condition for the enrichment score was generated, revealing no substantial differences across culture conditions for the underlying gene expression programs. **C)** The fidelity of HOTT tumor cell type to the GBM meta-atlas cell types was compared. Bubble plot, where the size of dots represents the cell number obtained by single cell transcriptome and the intensity of dots represents the predicted ID score, i.e., the similarity to GBM typical cell types. When compared to the first two columns representing primary tumors, the HOTT model demonstrates the best recapitulation of certain cell types including cycling, neuronal, vascular, and astrocyte. While microglia and MES were retained at relatively lower numbers and lower levels of similarity, they are still comparable to the scores of matched primary tumors from these samples. Furthermore, culture conditions without FBS favored cycling and radial glia cells, with M3noFBS+HEF showing the most dramatic effect. These results mirror previous discoveries in Fig 2, showing the robustness of our findings. Interestingly, OPC populations are also significantly favored in culture conditions without FBS. **D)** Differential gene expression analysis was performed between primary and transplanted tumors. Bar graphs of the top 10 most significant Reactome pathways were shown. Pathways related to both innate and adaptive immune systems were enriched in primary cells compared to transplanted cells, corroborating our observation of diminished immune cells in the HOTT system. **E)** When annotating non-neoplastic cells in HOTT, we observed high predicted.id.scores when using normal developing brain meta-atlas as reference (left panel). To annotate non-neoplastic cells from published GBM datasets, three references were applied. High predicted.id.score were observed when referenced to normal developing brain meta-atlas, but not normal adult or aged brain. This observation suggested that exposure to tumor might reactivate developmental program in surrounding brain cells in TME, highlighting the relevance of HOTT for the GBM microenvironment modeling.

In instances of neurological disease, transcriptomics programs relating to key features of functional biology can be altered in parallel cell types (Jourdon et al., 2023; Mathys et al., 2023) and we wished to explore whether individual media conditions impacted these transcriptional phenotypes. To perform this analysis, we leveraged two major strategies. First, we performed single-sample gene set enrichment analysis (ssGSEA) (Barbie et al., 2009; Subramanian et al., 2005), using the transcriptomic signatures of cell types in our meta-atlas as the reference gene sets. We then plotted the enrichment scores across all clusters and media conditions (Fig 3B), and that the distributions were similar across media conditions, suggesting transcriptional programs of parallel cell types across media conditions were largely unaltered. We additionally explored the predicted annotation scores when comparing our HOTT tumor cell types to our reference meta-atlas. We noted that HOTT better recapitulates certain cell types including cycling, neuronal, astrocyte, vascular and radial glial cells (Fig 3C). While mesenchymal cells are less well represented, they are comparable to the scores we generated for the matched primary tumors from these samples, indicating fidelity to their primary counterpart.

Given the reactivation of developmental programs in GBM, we also conducted additional module analysis to investigate the expression of recently described neurodevelopmental programs (Nano et al., 2023) across our HOTT media conditions compared to the primary tumor (Fig S3B). We observed similar module activity for most programs, again corroborating the fidelity of our model to the primary tumor context. Nuanced differences related to cell fate, metabolism and immune function were noted. We interrogated these differences further by performing differential gene expression analysis between the matched primary tumors and those in the HOTT system. We observed that immune function was upregulated in primary tumors (Fig 3D). This is expected given that we are unable to propagate immune cells in this system and that this is largely true of patient derived models (Akter et al., 2021). We observed an upregulation of programs related to translation, nervous system development and axon guidance in the HOTT system (Fig 3D), suggesting that the model promotes the expansion of the neurodevelopmental lineages. This may create a unique opportunity to study questions of cell fate specification in this system. Together, these data provided a nuanced understanding of which cell types are best recapitulated in the HOTT system, highlighting that although proportions of cells can vary based upon extrinsic environmental cues as modulated through the media, the heterogeneity and fidelity of these cell types was consistent across conditions.

Compared to other existing organoid models, HOTT uniquely enables the study of the direct interactions between the direct from patient tumor samples and a human TME. We sought to leverage this aspect of HOTT and first asked what cell types were present in the organoid microenvironment after exposure to the tumor. We annotated these non-tumor populations by projecting to cell types from the developing human brain meta-atlas (Fig 3E, SFig 4A). As expected, we observed high predicted scores and our annotations showed a variety of developmental cell types, consistent with the organoid as a developmental model. We then benchmarked this microenvironment to the patient TME by analyzing published datasets from which non-tumor cells were included in the single-cell analysis, a normal byproduct of resection. The same approach, i.e., projecting to a reference dataset, was also utilized to annotate these non-tumor cell types. Given these samples were from the adult human brain, we not only used our meta-atlas from the developing brain, but also a normal adult single-cell dataset (Hodge et al., 2019) as well as a study exploring the aging brain (Yang et al., 2022) as our reference. Fascinatingly, the normal cells obtained from adult tumor biopsies showed the highest similarity to the developing human brain, as opposed to the adult or aging brain (Fig 3E, SFig 4B). This highlighted the relevance of the organoid system for modeling the GBM microenvironment which, according to data collected here and our analysis of additional published datasets, deviates from the cell types found in the healthy adult brain (SFig 4B).

### HOTT uncovers bidirectional tumor - microenvironmental communication mediated by PTPRZ1

GBM cells are active participants in their own propagation and hijack resources from surrounding normal cells in order to promote their own growth and proliferation. HOTT offers a unique opportunity to study the effects of tumor cells on their microenvironment in a molecularly tractable, human-specific manner. With confidence in not only the tumor cell types in HOTT, but also the context in which they are grown, we asked whether anything about the organoids exposed to GBM differed compared to tumor naive organoids. To this end, we sorted out the GFP negative cells from the standard M3 condition, and across UMAP space observed segregation between the tumor naive and tumor transplanted organoid cells (Fig 4A). We noted that tumor-transplanted organoids showed a higher proportion of neuronally committed populations including intermediate progenitor cells, newborn neurons, and excitatory cells (Fig 4A). These changes in the organoid suggested communication occurs between the tumor and the organoid.

**Fig 4.**
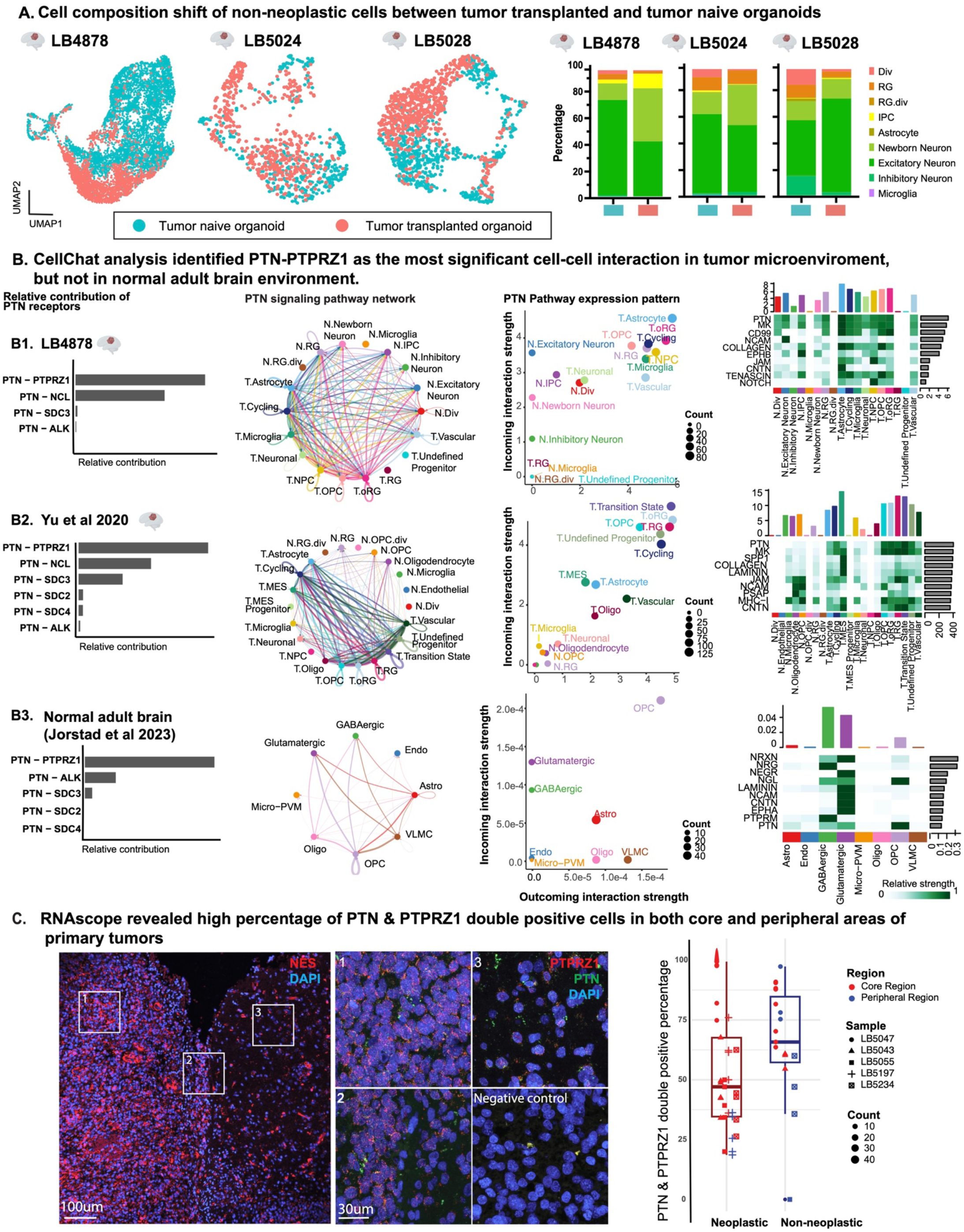
HOTT uncovers bidirectional tumor - microenvironmental communication mediated by PTPRZ1. **A)** When comparing non-neoplastic organoid cells from tumor-exposed organoids with batch-matched naïve organoid cells, we observed separation of these two samples in UMAP space across all three primary GBM experiments (left panel). Cell composition analysis (right panel) revealed a persistent shift from dividing cells and progenitors towards neuronally committed populations after exposure to GBM, suggesting that GBM may promote the differentiation of surrounding normal cells. **B)** Cell-cell communication in the tumor microenvironment was assessed using CellChat (Jin et al., 2021), a computational tool that enables prediction of cell interaction based on receptor - ligand relationships in a cell type specific manner. B1) In the HOTT system, represented by LB4878 in the M3 culture condition, PTN was identified as the most significant signaling pathway (right column) with PTPRZ1 as its major receptor (left column). The PTN signaling pathway network was shown in a circle plot. Dominant sources of this signaling as both sources and targets, included neoplastic dividing cells, progenitors and radial glia, as shown in the 2D scatter plot. B2) When leveraging the same strategies to compare the neoplastic and non-neoplastic cells in a published GBM dataset (Yu et al), by comparing PTN - PTPRZ1 was again identified as the top signaling between the tumor and the microenvironment. B3) However, when a similar analysis was conducted with a normal adult human brain dataset, this pathway did not emerge as a primary mediator of cell-cell interactions. Note the significantly lower PTN interaction strength observed in the 2D scatter plots within the normal adult brain as compared to the GBM TME. The interaction strength across analyses demonstrates higher scores in the neoplastic cell types in tumor samples compared to normal cell types. **C)** RNAscope revealed high percentages of PTN and PTPRZ1 double positive cells in both core and peripheral areas of primary tumors. FFPE sections were derived from surgical samples removed from the tumor core (core region) and the tumor boundary (peripheral region). Neoplastic regions were identified by post-staining with NES on contiguous sections, as well as by assessing cell density based on DAPI nuclei staining. NES staining on the right is autofluorescence, staining on the left with cellular co-localization is true staining. Almost all the cells are PTN and PTPRZ1 double positive, mirroring the equal incoming and outcoming interaction strength in Fig 2B 2D scatter plots. The observation of PTPRZ1 and PTN positive cells found in both the neoplastic and non-neoplastic areas of the tumor in both the core and peripheral regions supports the role of PTN-PTPRZ1 in potentially mediating tumor migration.

Thus, we investigated potential key signaling pathways that could act as mediators between tumor and their surrounding non-neoplastic cells by inferring ligand-receptor interactions with CellChat (Jin et al., 2021) and observed that PTN-PTPRZ1 signaling was the most significant signaling pathway upregulated in the HOTT tumors (Fig 4B, SFig 5). This was intriguing since this signaling pathway that has been implicated in tumor migration (Qin et al., 2017), including through GBM outer radial glia (Bhaduri et al., 2020b). Because the organoid is a developmental model and both PTN and PTPRZ1 are widely expressed during development (Pollen et al., 2015), we were wary of a model specific artifact influencing these results. Further interrogation of datasets from patients containing non-neoplastic microenvironmental cells also showed higher expression of PTPRZ1 and PTN in the TME compared to the normal adult brain (Krawczyk et al., 2022; Wang et al., 2022), giving us confidence in our system to identify novel tumor - TME interactions.

In the context of patient tumors (Yu et al., 2020), we again observed that PTN - PTPRZ1 signaling between the tumor and the microenvironment topped the predicted list of interaction pathways (Fig 4B). Notably, we observed that in tumors PTN and PTPRZ1 were highly expressed in all tumor cell types, with overall lower expression in the normal peritumor cells (Fig 4B). A similar trend was observed with other published datasets containing primary GBM tumors and normal cells (Bhaduri et al., 2020b; Darmanis et al., 2017) (SFig 6). However, when a similar analysis was performed with the healthy adult human brain (Jorstad et al., 2023), this pathway was not a top pathway mediating cell - cell interactions (Fig 4B), and overall interaction strength was observed to be substantially lower than in the HOTT and GBM patient datasets (Fig 4B, SFig 6). This reinforces the notion that PTPRZ1 communication may result from the reactivation of developmental programs previously observed in GBM. The TME activation of PTPRZ1 signaling is specific to the tumor and peritumor environment, potentially mediating a crucial axis of tumor – normal cell communication (Fig 4B). These findings were consistent across samples, datasets, and analyses, including additional HOTT samples, published datasets, and transcriptional profiles of the adult healthy human brain (SFig 5 - 6).

We sought to validate these findings in newly resected patient tumor samples. Patient tumor samples were surgically removed from the tumor core (core region) and from the tumor boundary (peripheral region) as defined by a recently described amine chemical exchange saturation transfer echo planar imaging (CEST-EPI) approach (Patel et al., 2024). Because of heterogeneity in the spatial distribution of GBM, both resections contained neoplastic and non-neoplastic cells, which is consistent with what has been described in published studies (Wang et al., 2022). Thus, we sought to identify the neoplastic cells in our *in situ* analysis by post-staining the analyzed samples with Nestin (NES). Neoplastic cells were identified based upon the distribution of this staining as well as differences in cell density. Our RNAscope analysis revealed that PTPRZ1 and PTN were found in both the neoplastic and non-neoplastic cells of the tumor in both the core and peripheral regions (Fig 4C). When analyzing these data, we noted that almost all cells were double positive for both PTN and PTPRZ1, suggesting a joint role in tumors. Additionally, the fact that PTN and PTPRZ1 are in the peripheral surgical sample supports their role in potentially mediating tumor migration, and these data strongly suggested a role for novel signaling interactions between the tumor and the TME.

### Microenvironmental PTPRZ1 has an unexpected role in mitigating tumor migration

PTPRZ1 is primarily known for its cell intrinsic role in promoting tumor stemness and invasion. Notably, PTPRZ1 has long been described as a receptor tyrosine phosphatase (Xia et al., 2019) that maintains glioma stemness and drives proliferation (Fujikawa et al., 2017). Additionally, *in vivo* work showed that PTN ligand binding to PTPRZ1 resulted in increased tumor invasion through Rho/ROCK signaling (Qin et al., 2017). In our previous work, we identified PTPRZ1 as a marker for GBM outer radial glia and showed that PTPRZ1 was required for the outer radial glia cell jump and divide cell behavior known as mitotic somal translocation (Bhaduri et al., 2020b; Ostrem et al., 2017). Previous work has not identified a role for PTPRZ1 in communicating with the TME, though PTN has been described to be secreted by microenvironmental macrophages (Shi et al., 2017) and has also been characterized as a chemoattractant for tumor cells (Qin et al., 2017). Meanwhile, our data suggests co-localization of PTN and PTPRZ1 in both the tumor and in the peritumor environment using both single-cell analysis and RNAScope characterization of patient tumor samples.

This led us to hypothesize that PTPRZ1 may be involved in tumor migration through a cell extrinsic role via tumor – TME communication. To explore how migration might be impacted by both intrinsic and environmental PTPRZ1 expression, we devised a novel migration assay in which we transplanted dissociated patient derived gliomaspheres to organoids using our hanging drop method. We used gliomaspheres for this experiment because they enabled us to query both tumor intrinsic phenotypes in culture and to examine if these phenotypes were the same or different after introduction to HOTT. After 2 days in culture, we generated fusion organoids between transplanted and gliomasphere naive organoids. Each fusion served as a migration experiment whereby tumor cells from the transplanted side had an opportunity to migrate across the fusion boundary. We evaluated migration using both cell sorting and immunostaining (Fig 5A). For the sorting experiments, we severed the fusions at the junction point, identifying the original transplanted side as the one with GFP expressing cells fully surrounding the organoid, though migration to the other side could be visualized when looking at the live fusion experiment (Fig 5B). GFP positive cells could then be sorted from each side of the fusion, enabling a quantitative metric of how much migration was observed.

**Fig 5.**
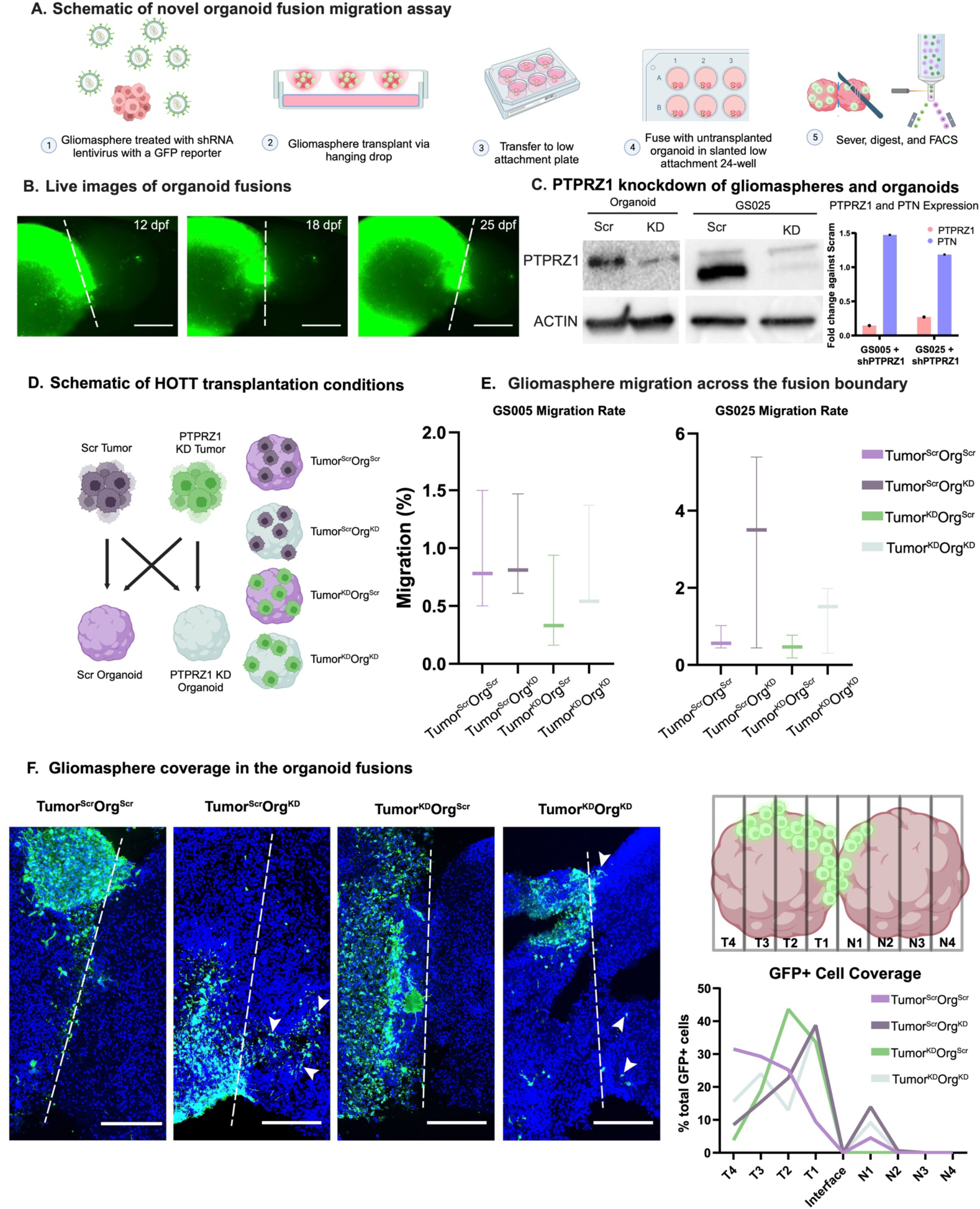
Environmental PTPRZ1 knockdown increases glioma migration. Workflow of a novel organoid fusion migration assay. Gliomaspheres were treated with a shRNA lentivirus that contains a scramble or PTPRZ1-targeting sequence with a GFP reporter prior to transplantation. After transplantation using the hanging drop method, the HOTTs were allowed to fuse with a tumor-naive organoid and cultured for 3 weeks. At time of harvest, the fusions were severed with a scalpel and digested for FACS analysis. **A)** Images of the HOTT fusions at 12, 18 and 25 days post fusion (dpf). GFP signal represents engrafted GS. White dotted line represents the fusion boundary. Scale bar = 750 μm. **B)** Western blot analysis of organoid and GS used for transplantation (left and middle). PTPRZ1 protein levels significantly decreased following shRNA lentivirus treatment. Real-time PCR of gliomaspheres show PTPRZ1 knockdown after shRNA was introduced, while PTN expression was unchanged (right). **C)** Schematic representation of the various transplant combinations. GS cells were treated with either a scramble shRNA or PTPRZ1 targeting shRNA and transplanted onto scramble or PTPRZ1 knockdown organoids, resulting in 4 experimental conditions. **D)** Box-and-whisker plots of migration rate for two patient-derived gliomasphere lines, GS005 and GS025. The migration rate was calculated by dividing the number of GFP+ cells on the tumor-naive side with the total number of GFP+ cells from both sides (n=3 replicates per line). Under identical tumor PTPRZ1 KD status, organoid PTPRZ1 KD resulted in increased GS migration towards the tumor-naive side. **E)** Immunostaining of the organoid fusions at the fusion boundary (left). Migration of GFP labeled GS cells was observed. Under identical tumor PTPRZ1 KD status, organoid PTPRZ1 KD resulted in increased GS migration towards the tumor-naive side (example labeled with white arrowheads). Scale bar = 250 μm. Schematic of the binning strategy employed to quantify the distribution of GS cells was shown on the top right. The organoid fusion was divided into eight equal bins labeled T1-4 (transplant) and N1-4 (naive) which stem from the fusion boundary. The amount of GFP+ cells in each bin was quantified and plotted as a ridge graph as the percentage of total GFP cells present in each bin (bottom right).

For these migration studies, we leveraged the versatility of the HOTT system in enabling us to knockdown environmental PTPRZ1, thus evaluating whether TME PTPRZ1 contributes to the migration phenotype. Specifically, we performed knock-down of PTPRZ1 in organoid cells using a small hairpin RNA (shRNA) before transplantation with gliomasphere cells, themselves treated with either a scrambled or PTPRZ1-targeting shRNA (Fig. 5D). Analogous transplantations were conducted in scrambled shRNA-expression organoids. We performed tumor and/or organoid knockdown for PTPRZ1. Migration was then evaluated (Fig 5D); across both the organoid and the gliomaspheres we observed moderate PTPRZ1 knockdown (Fig 5C, SFig 7). Importantly, knockdown of PTPRZ1 had no impact on PTN expression, ensuring we were studying PTPRZ1, PTN independent, effects (Fig 5C).

From the FACS quantification, we validated previous studies highlighting that tumor knockdown of PTPRZ1 decreased overall migration rate (Bhaduri et al., 2020b; Qin et al., 2017; Shi et al., 2017). However, environmental PTPRZ1 knockdown increased tumor migration: tumors transplanted into organoids where PTPRZ1 was knocked-down migrated more efficiently than those transplanted in scrambled shRNA-expressing organoids. This was true both in the context of tumor scrambled hairpin and tumor PTPRZ1 knockdown (Fig 5E). This was further validated using an orthogonal measurement of migration where immunofluorescence for GFP labeled gliomasphere cells was quantified on either side of the fusion border (Fig 5F). Again, knockdown of PTPRZ1 in the TME corresponded to greater tumor migration. These data contradict our expectations about PTPRZ1; we assumed based upon previous studies that PTPRZ1 knockdown, even in the TME, would reduce migration. However, our data suggest a distinct role for how peritumoral PTPRZ1 may be inhibiting tumor migration, and when removed alters aspects of tumor – TME communication that drives behavioral changes.

### Microenvironmental manipulation of PTPRZ1 alters tumor cell fate through a catalytically independent mechanism

Previous work has described the role of intrinsic PTPRZ1 in maintaining tumor stemness and suppressing oligodendrocyte precursor cells differentiation (Fujikawa et al., 2017). To explore if this cell - cell communication extended to properties of cell fate specification, we performed immunostaining for the major cell types identified in our single-cell analysis and quantified our observations. Again, we were surprised to see that environmental knockdown of PTPRZ1 modulated changes in tumor cell types compared to the control conditions, even without manipulation of the tumor itself. This implies a degree of cell nonautonomous tumor - microenvironmental communication. For example, environmental knockdown of PTPRZ1, without manipulation of the tumor directly, drives reduction of the tumor mesenchymal fraction as marked by HMOX1 and of the tumor neuronal fraction as marked by MEF2C (Fig 6A-B). Additionally, TME knockdown of PTPRZ1, again without direct manipulation of the tumor itself, increased tumor astrocyte populations as marked by GFAP (Fig 6C-D); the reduction of neuronal populations and increase in immature astrocytes suggests a shift towards a more stem-like identity. When targeting the tumor directly, tumor knockdown of PTPRZ1 increased both tumor and organoid CTIP2, a marker of mature deep layer neurons, including when the microenvironment was not directly manipulated, indicating additional tumor and environmental differentiation which is consistent with previous PTPRZ1 literature. Interestingly, the increase in CTIP2 in the organoid is also spatially located near the transplantation, suggesting a local effect consistent with our hypothesis that these cell fate changes are mediated by cell - cell communication through PTPRZ1 (Fig 6C). In support of this hypothesis, we observed that the cell fate changes that occurred upon transplantation of the gliomaspheres were not the same effects observed when looking at gliomasphere cell type composition on its own (SFig 7 D-E), further emphasizing that the microenvironment can change tumor intrinsic properties of cell fate transition.

**Fig 6.**
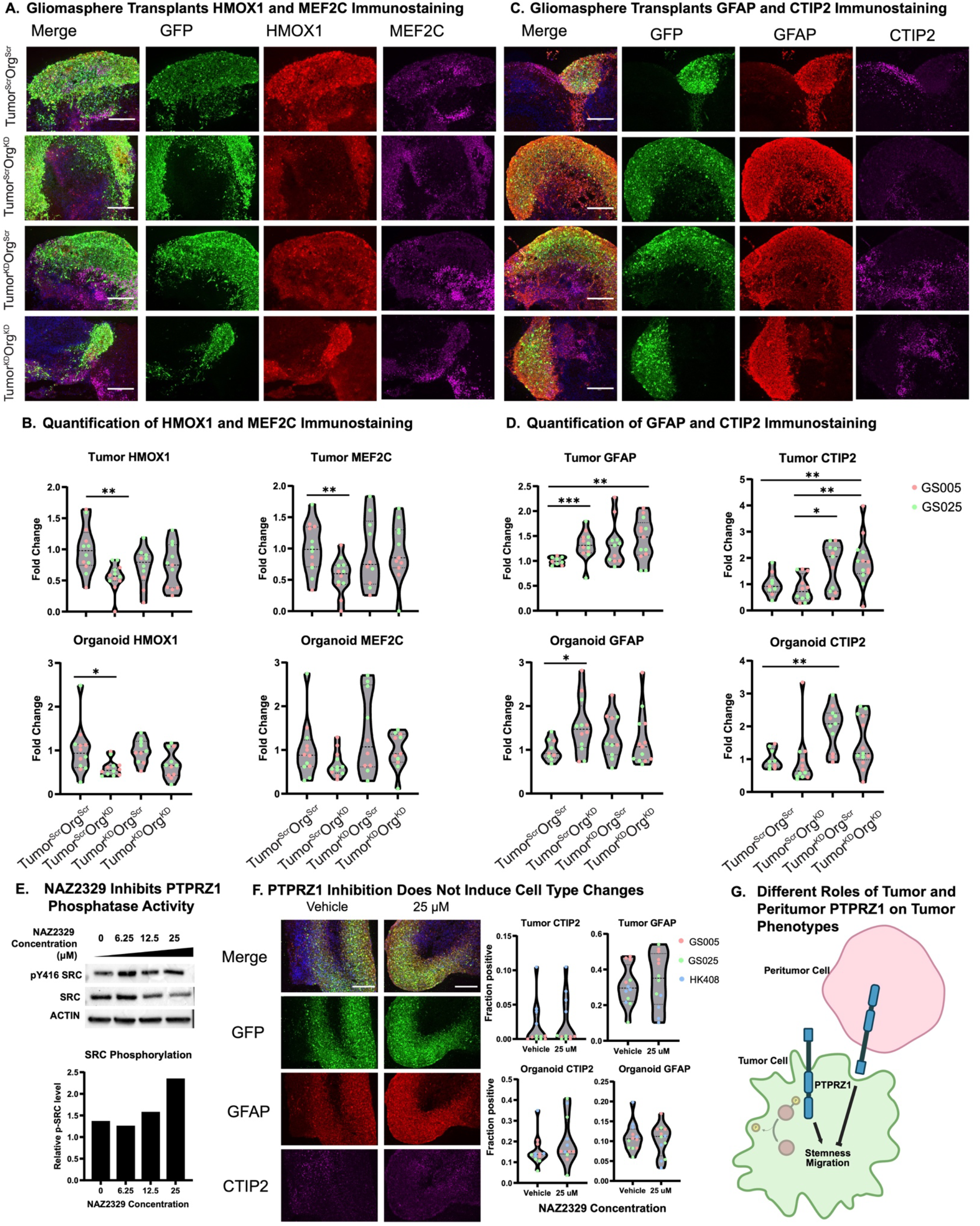
Environmental knockdown of PTPRZ1 alters tumor cell fate. **A)** Effects of tumor or environmental PTPRZ1 KD on cell fate were assessed with immunostaining of mesenchymal marker HMOX1 and neuronal marker MEF2C. Organoid KD of PTPRZ1 significantly reduces the expression of tumor HMOX1 and MEF2C. Moreover, organoid HMOX1 expression was also reduced with organoid PTPRZ1 KD, suggesting a role for PTPRZ1 in driving mesenchymal transition both intrinsically and through cell-cell communication. **B)** Violin plots depicting the quantified tumor and organoid expression of HMOX1 and MEF2C. Fold change of each experimental condition was calculated based on the average of all the data points in Tumor^Scr^Org^Scr^. Tumor and organoid expression was based on the ratio of GFP+/Signal of interest+ over total GFP+ cells and GFP-/Signal of interest+ over total GFP-cells, respectively. P-value determined by two-tailed Mann-Whitney test, *p ≤ 0.05, **p ≤ 0.01, ***p ≤ 0.001. Experiments were conducted in two patient-derived gliomasphere lines (GS005 and GS025) with a minimum of 3 but up to 6 replicates per line. **C)** Effects of tumor or environmental PTPRZ1 KD on cell fate were assessed with immunostaining of astrocyte marker GFAP and deep layer neuronal marker CTIP2. Organoid KD of PTPRZ1 increased tumor GFAP and organoid GFAP expression, suggesting a role for PTPRZ1 in regulating astrocyte differentiation. In addition, under the context of organoid PTPRZ1 KD, tumor PTPRZ1 KD significantly increased tumor CTIP2 expression. On the other hand, tumor PTPRZ1 KD increased organoid CTIP2 expression, suggesting a role for neuronal fate regulation via tumor-normal PTPRZ1 signaling. **D)** Violin plots depicting the quantified tumor and organoid expression of GFAP and CTIP2. Fold change and statistical significance were calculated as described in Fig. 6B. **E)** Western blot analysis of NAZ2329 treated GS cells at concentrations of 0 (Vehicle), 6.25, 12.5 and 25 uM. Total SRC and pY416 SRC levels were examined and the ratio of pSRC to SRC levels increased in a dose-dependent manner, validating the blockage of PTPRZ1 phosphatase activity. Actin was used as a loading control. 30 ug of protein was loaded in each lane. **F)** Effects of cell fate driven by PTPRZ1 catalytic inhibition was investigated by immunostaining of astrocyte marker GFAP and neuronal marker CTIP2 (left). The organoid transplants were treated with a DMSO vehicle or 25 μM of NAZ2329. Violin plots showing quantification of GFAP and CTIP2 expression (right). P-value determined by two-tailed Mann-Whitney test, *p ≤ 0.05. Experiment was conducted with three GS lines with four replicates per line. **G)** Graphic model illustrating the different roles of tumor derived or peritumor derived PTPRZ1 on tumor stemness and migration. Intrinsic tumor PTPRZ1 drives tumor stemness and migration, whereas microenvironmental PTPRZ1 suppresses both. This mechanism is independent of PTPRZ1 phosphatase activity.

As a receptor tyrosine phosphatase, PTPRZ1 is most well described to function through its catalytic activity by which it removes phosphate groups from downstream targets including Src (Xia et al., 2019). To test whether these changes in cell fate are mediated by PTPRZ1 catalytic activity, we utilized a small molecule inhibitor, NAZ2329, that inhibits the catalytic activity of PTPRZ1 by binding to its active site in the D1 domain (Fujikawa et al., 2017). At 25 μM, NAZ2329 inhibition increased the accumulation of phospho-SRC as expected (Fig 6E). Since the small molecule was added to the culture media, PTPRZ1 inhibition occurred on both gliomasphere and organoid cells, and so was most closely resembling the tumor and organoid double knockdown condition. Notably, the increase in tumor GFAP and CTIP2 population observed in this double knockdown was lost in the NAZ2329-treated condition (Fig 6F), suggesting the cell type changes are not mediated by PTPRZ1 catalytic activity.

Overall, we observed that TME PTPRZ1 knockdown is increases tumor immature populations, while tumor knockdown augments mature populations. This contradictory role is further consistent with our observations of migratory phenotypes, again, where the environmental PTPRZ1 is a block on migration. This suggests a push - pull model of tumor intrinsic and extrinsic roles for PTPRZ1 signaling (Fig 6G), and specifically highlights the need to study GBM signaling pathways and biology in the context of a microenvironment as we achieve with our HOTT system.

## Discussion

Effective models of GBM that recapitulate the heterogeneity of human patient tumors and their complex interactions with the TME are a crucial tool in the field of neuro-oncology to overcome existing barriers in designing effective treatments for this devastating cancer. Advances in the development of organoid models have opened up vast opportunities in the study of GBM and addressed significant limitations of existing models of 2D cell culture and mouse models. The optimized HOTT system that we present here and benchmark against both the patient tumor as well as the patient microenvironment allows for the study of tumor-TME crosstalk in a human context. Notably, we observed tumor fidelity in both gene expression and cell type composition to the tumor of origin in our HOTT system. Furthermore, the gene expression of each cell type remained consistent across media conditions, including FBS-containing culture conditions, suggesting that the existence of normal brain cells around the tumor could be an essential driver in maintaining cell type identity.

Here, we leveraged HOTT to explore tumor - TME interactions, inadvertently stumbling upon a crucial role for PTPRZ1 in enabling both the tumor and the TME to reciprocally influence one another. The ability for a tumor to mold the microenvironment is described in other systems (de Visser and Joyce, 2023), and in the context of GBM is largely contextualized in terms of the immunosuppressive forces the tumor exerts on the immune populations in the environment (Pombo Antunes et al., 2020). Recent efforts in the field of cancer neuroscience have begun to highlight mechanisms by which the TME can promote tumor growth and integrate into the electrophysiological networks of the brain. Our finding that PTPRZ1, a receptor tyrosine phosphatase, can promote cell - cell communication adds a branch to the ways in which GBM functionally communicates with its TME.

Our HOTT migration assay is a novel tool to explore tumor migration in the context of human TME. Commonly used migratory assays in two-dimensions and in mice either entirely lack a microenvironment or rely on non-human contexts. An advantage of HOTT is that it is molecularly tractable, making knockdown experiments in this system substantially more feasible than generating a knockout mouse when interrogating an individual candidate gene. Using this system, we were also able to identify cell fate shifts mediated by reciprocal knockdown; i.e. tumor knockdown changed the properties of the TME and TME knockdown altered cell fate specification in the tumor. This clearly suggests a non-cell autonomous role for PTPRZ1 signaling, independent of its catalytic activity, that has not been documented in the existing literature.

Our work here shows that PTPRZ1 plays an unexpectedly significant role in driving tumor – TME communication. Altogether, the HOTT system, our analysis of published patient data, and our RNAscope analysis of GBM margins highlights that PTPRZ1 is present both in the tumor and the peritumor environment. We and others have previously described a role for PTPRZ1 in promoting tumor stemness and migration (Bhaduri et al., 2020b; Fujikawa et al., 2017; Mikelis et al., 2009; Shi et al., 2017), whereby PTPRZ1 is required for these pro-tumorigenic properties. This aspect of PTPRZ1 has proven puzzling; in various models it has been shown that not only is PTPRZ1 required for migration and glioma stemness, and that this is a function of its catalytic activity (Fujikawa et al., 2016; Fujikawa et al., 2017). However, PTPRZ1 is a phosphatase, removing phosphate groups from downstream pathways such as Src and beta catenin, where the accumulation of phosphorylation is pro-tumorigenic (Meng et al., 2000). Interestingly, previous work has also suggested that PTPRZ1 is most tumorigenic when bound by its ligand, PTN, which interestingly results in the dimerization of the protein, inactivating its catalytic activity.

PTPRZ1 is not widely expressed in the adult human brain (Jorstad et al., 2023), but it is upregulated as we and others note across GBM cell types (Xia et al., 2019). Why, then, would glioblastoma cells upregulate PTPRZ1, just to inactivate it? If PTPRZ1 is playing a substantial role in cell - cell communication as we suggest here, this cell extrinsic role explains both why catalytically inactive, but highly expressed, PTPRZ1 could be highly functional for glioblastoma cells. We support this hypothesis by our inhibitor-based experiments in which we show these impacts upon cell fate specification are dependent on PTPRZ1 expression but not catalytic activity. Future efforts in the HOTT system and in other models will be required to further elucidate whether common downstream pathways are activated by cell extrinsic PTPRZ1 communication as are invoked in traditional PTPRZ1 signaling, and whether the cell – cell communication roles of PTPRZ1 are modulated by PTN or other ligands that may exist in the tumor milieu.

Importantly, these findings change our interpretation of PTPRZ1 as a high value therapeutic target; our previous work, work of others, and our own data in this study highlight that when tumor cells are investigated alone, PTPRZ1 knockdown is a valuable strategy to decrease migration and promote tumor differentiation. However, it is unlikely that most drugs would specifically target the tumor only, and our observations of TME knockdown of PTPRZ1 show antagonist effects. Though the eventual impact on patient tumors cannot be extrapolated, there is a risk that the apparent inhibitory role on tumor progression that the TME plays by expressing PTPRZ1 would be lost with therapeutics targeting PTPRZ1, mitigating the effect of a drug or even worsening overall progression. Thus, any therapeutic approaches seeking to target PTPRZ1 must ensure impressive tumor specificity, or should use alternative approaches to target this important signaling protein at the nexus of migration and cell fate in GBM.

A fundamental result of this study is that therapeutic efficacy in GBM will eventually rely on treating the whole brain, and thus, the tumor plus its complex TME. The field is aware of this complexity which underlies numerous drug trials that are less efficacious in patients compared to other models (Bagley et al., 2022). Systems like HOTT enable scalable experimental investigation of direct from patient tumor cells in a relevant microenvironmental context. Eventually, one could imagine that future efforts would directly target the microenvironment, which provides the valuable “soil” in which GBM thrives. However, HOTT too is currently limited in its ability to recapitulate the full microenvironment as it lacks a functional immune system. We have observed that when tumors are transplanted into HOTT, few to no immune cells survive after 3-4 weeks culture, and future efforts should focus on extending this capability of the model. At present, however, HOTT provides an essential additional layer of complexity, modeling important aspects of the human GBM TME, and providing novel insights into the ways GBM and the TME functionally interact.

## Data Availability

Data from this study are available for download and interactive analysis via our Cell Browser at: https://hott.cells.ucsc.edu.

## Acknowledgments

We would like to thank the members of the Bhaduri Lab for their insightful advice and comments on the study. We would like to thank the Broad Stem Cell Research Center Flow Cytometry core for their help in isolating cells for this project, and Charina Julian (UCSF) for help with running sequencing. We would like to thank Sergey Mareninov and others at the Brain Tumor Translational Research Core at UCLA for enabling tumor sample acquisition. This study was generously funded by funding from to AB from: Swim Across America, Jonsson Comprehensive Cancer Center, Sloan Research Fellowship from the Alfred P. Sloan Foundation, NCI P50CA211015 including a Career Enhancement Program Award, The Sontag Foundation (Distinguished Scholar Award), V Scholar Award from The V Foundation, The Uncle Kory Foundation, The American Cancer Society (CSCC-Team-23-980262-01-CSCC), The Margaret Early Medical Research Trust, The Rose Hills Foundation and the Broad Stem Cell Research Center. Additional funding was obtained from NIH T32GM145388 (RLK). Mostly, we would like to thank the patients who generously donated their tumor tissue and entrusted us to make progress on this devastating disease.

## Author Contributions

Study conceptualization was performed by AB, WG, and DAN. Data curation was performed by WG, EF, PRN. Formal analysis was performed by WG, RLK, CY, EF, PRN, DJA, JS, AB. Funding acquisition was performed by AB and RLK. Investigation was performed by WG, RLK, CY, EF, PRN, DJA, ML, JYI, CT, HAT, JS, and KP. Methodology was developed by WG, RLK, EF, PRN, DJA, KP, DAN and AB. Project administration and supervision was performed by AB. Resources were generated by WG, RLK, and KP. The original draft was generated by AB, RLK and WG with editing and revision from all authors.

## Methods

### Stem Cell Culture and Maintenance

Human embryonic stem cells (hESCs) and human induced pluripotent stem cells (hiPSCs) were cultured consistent with other studies (Bhaduri et al., 2020a; Pollen et al., 2019; Velasco et al., 2019). Multiple hESC/hiPSC lines derived from UCLA and CIRM were utilized in this study. In brief, stem cells were cultured on Matrigel (FisherScientific, 354230) coated 6-well plates in mTeSR plus supplemented with 10% mTeSR plus Supplement (Stem Cell Technology, Cat 100-0276), and 1X Penicillin/Streptomycin or Primocin (FisherScientific NC9141851). Media changes were performed every other day, and passaging was conducted when cells reached over 75% confluence. During passaging, ReLeSR (Cat 5872) was applied at room temperature for 1 minute, aspirated, and the stem cells were left at 37°C for 5 minutes before fresh stem cell media were added to dissociate them into smaller clusters without reducing them to single cells. These smaller clusters were evenly distributed into new Matrigel-coated plates at a 1:4 or 1:6 ratio. For freezing, the same procedure as in stem cell passaging was followed, except in the final step, 1 ml of mFreSR (Cat 5855) per well in a 6-well plate was used to resuspend the stem cells. The resuspended cells were then transferred to cryovials for storage at -80°C for 24-48 hours before being transferred to a liquid nitrogen tank for long-term storage.

### Cortical Organoid Generation

Cortical organoids were generated using an adapted version of the method described first in Kadoshima et al 2013, similar to other studies (Bhaduri et al., 2020a; Bhaduri et al., 2020b; Kadoshima et al., 2013; Velasco et al., 2019). Briefly, 1ml Accutase (Sigma A6964) was added to each well in a 6-well plate with human embryonic stem cells (hESCs) having over 75% confluence. After a 37°C incubation for 5 minutes, 1 mL of Media 1 (M1) was added to each well. M1 consists of GMEM (LifeTech 11710-035), 20% KnockOut Serum (LifeTech 10828-028), 0.1 mM β-mercaptoethanol(Sigma 21985023), 1X Non-Essential Amino Acids (NEAA), 1X Sodium Pyruvate, and 1X Penicillin/streptomycin or Primocin (FisherScientific NC9141851). The cells were scraped out and transferred to 15 mL tubes. After centrifugation at 300 x g for 5 minutes, cell pellets were resuspended into a single-cell suspension with 1 mL of M1 supplemented with three small molecules: Rock inhibitor Y27632 (20 μM), TGF-β inhibitor SB431542 (5 μM), and Wnt signaling inhibitor IWR1-endo (3 μM). One million cells in 10 mL of M1 supplemented with the small molecules were evenly distributed into a 96-well V-shaped low attachment plate (S.Bio MS-9096VZ). Cells were allowed to aggregate undisturbed for 72 hours. On the 3rd day, the first media change was performed by replacing 50 μL with 100 μL of fresh M1 containing the small molecules to avoid disturbing the nascent organoids. Subsequently, 100% media changes were performed every other day until Day 7 when the Rock inhibitor was removed from the recipe. At 18 days, the organoids were transferred to a low attachment 6-well plate (Fisher Scientific 07-200-601). Subsequent media changes were performed every other day. From Day 18 to Day 35, Media 2 (M2) was used, which consists of DMEM/F-12 with Glutamax (Lifetech 10565-018), 1X N-2 supplement (Lifetech 17502-048), 1X Lipid Concentrate (Lifetech 11905-031), and 1X Penicillin/streptomycin or Primocin. Starting from Day 35, Media 3 (M3) was applied, which includes DMEM/F-12 with Glutamax, 1X N-2 supplement, 1X Lipid Concentrate, 1X Penicillin/streptomycin or Primocin, 10% Fetal Bovine Serum (GIBCO 10082147), 5 μg/mL Heparin (Sigma H3149), and 0.5% Growth factor reduced Matrigel (Corning 354230). Throughout the culture period, live images were periodically taken to monitor organoid growth, and immunostaining was performed at Week 5 and Week 8 to confirm normal neuronal differentiation.

### Media Conditions Used for Experiments

Five different culture conditions were used in this study:

1. M3 served as a based media for comparison.
2. M3+B27: M3 media supplemented with 10% B27 (ThermoFisher Scientific 12587010)
3. M3noFBS: M3 media without FBS supplemented with 10% of DMEM/F-12 with Glutamax (Lifetech 10565-018).
4. M3noFBS+HEF: M3noFBS supplemented with 20 ug/mL Hu EGF (Life Technologies PHG0313), 8 ug/mL Hu FGF-b (Life Technologies cat. # PHG0263) and 2 mg/mL Heparin (Sigma-Aldrich H3149) as their final concentration.
5. M3noFBS+PDGF: M3noFBS supplemented with 50ng/ml PDGFAA (GIBCO PHG0035) as its final concentration.

### Tumor Dissociation and Viral Transduction

Primary tumors were obtained from Ronald Reagan Hospital at UCLA with appropriate consent forms for genomic analysis covered by Institutional Review Board IRB numbers 10-000655 and 21-000108. Tumors are cut into smaller pieces with a sterile scalpel blade. The material was then transferred to a 5 mL microcentrifuge tube containing 2.5 mL Papain supplemented with 125 uL DNase (Worthington LK003150) and incubated for 30-45 minutes in a 37°C incubator. After the initial 10 minutes of incubation, the tube was taken out and shaken vigorously for 10 seconds by hand to aid dissociation. This process was repeated after every 5 minutes of incubation thereafter. The material was further broken down with trituration before it was pelleted at 300g for 5 minutes. The pellet was then resuspended in warm media #3 and filtered with a 40 μM mesh. The cells were then infected with a GFP virus (SignaGen SL100268) at a concentration of 1 uL / 1 million cells supplemented with 1:1000 Polybrene Infection Reagent (Millipore Sigma TR-1003-G) for 90 minutes on a rotator in the 37°C incubator. The cells were subsequently washed three times with warm media before proceeding to organoid transplantation.

### Tumor Transplantation

After lentiviral infection, cells were counted after being resuspended in 1ml of warm media 3. Depending on the total tumor recovered and experimental design, the whole single cell suspension was distributed into several Eppendorf tubes, one tube for each culture condition. Following a 5 minute spin at 300g, the cell pellet was resuspended in the final volume of each culture media. 500K-1M tumor cells were transplanted onto one organoid in a 10-15ul droplet of corresponding culture media with the following methods. Hanging drop method: To set up the hanging drop tumor transplantation, 8-12-week-old human cortical organoids were transferred onto the lid of a 10 cm dish using wide-bore 1000ul tips. After removing extra media, 10-15ul tumor single cell suspension was added on top of each organoid. 6 organoids of the same culture condition were placed on one lid. Then the lid was carefully flipped onto the 10 cm dish with 10ml base culture media to prevent the media evaporating during the hanging drop stage. These hanging drop co-cultures were 50 maintained for 12 hours in the 37C incubator, before they are moved back to ultra-low attachment 6 well plate (Fisher Scientific 07-200-601) with corresponding culture media. Tumor cells usually surround the organoids and migrate inside after a few days. Such cultures last for about 4 weeks with 3 times media change each week before being harvested for analysis.

To optimize GBM transplantation, we employed another two methods, i.e., Insert and V-bottom approaches for a head-to-head comparison. For insert transplantation, 8–12-week-old organoids were positioned at the air– liquid interface on Millicell (Millipore) inserts to restrict movement. A 10-15 μl tumor single cell suspension was gently pipetted onto the semi-dry organoids, allowing integration for 30 minutes at 37°C. Subsequently, organoids were carefully lifted from the inserts by increasing the medium volume by 3 ml. After 2 days, organoids were transferred to an ultra-low attachment 6-well plate without inserts, maintaining appropriate culture media. For V-bottom transplantation, 8–12-week-old organoids were placed into V-bottom low attachment 96-well plates, with one organoid per well and 100 μl of appropriate media. A 10-15 μl tumor single cell suspension was slowly pipetted onto each organoid. After 2 days, organoids were transferred to an ultra-low attachment 6-well plate with suitable culture media.

### FACS Isolation of Tumor Cells

Tumor-transplanted organoids were transferred to a 1.7 mL Eppendorf tube containing 1 mL Papain supplemented with 50 uL DNase (Worthington LK003150) and incubated for 30-45 minutes in a 37°C incubator. After the initial 10 minutes of incubation, the tube was taken out and shaken vigorously for 10 seconds by hand to aid dissociation. This process was repeated after every 5 minutes of incubation thereafter. The material was further broken down with trituration before it was pelleted at 300g for 5 minutes. The pellet was then resuspended in cold FACS buffer (PBS + 2% FBS), filtered with a 40 μM mesh, and stained with DAPI to identify non-viable cells. DAPI-/GFP+ tumor cells and DAPI-/GFP-normal cells were sorted and collected for downstream single-cell capture.

FACS was performed on either a BioRad S3e or BD Biosciences FACSAria II Cell Sorter. A 100 μM nozzle was used for sorting. All sorts were performed with a purity sort precision at a slow speed. Gates were set to remove debris, eliminate doublets, and DAPI staining was used to exclude dead cells. Gates for GFP positivity were set using control, batch matched organoids without transplantation.

### Single-cell Capture and Sequencing

Single-cells from FACS or from dissociated tumor cells were captured using the 10X v3.1 3’ capture protocol and the 10X Chromium Machine. 10,000 cells were targeted for capture, or if the retrieved number after sorting was less than 20,000, all cells were used for capture. Capture proceeded as detailed in the user manual and manufacturer guidelines were used for library preparation. Sequencing was performed on an Illumina NovaSeq 6000.

### Single-nuclei Capture

Single-nuclei sequencing was conducted on all samples to capture the full spectrum of patient tumor cell types. Nuclei extractions were carried out from frozen tissue specimens obtained from the UCLA Brain Tumor Translational Resource (BTTR) or from flash-frozen samples preserved prior to dissociation. Frozen tissue samples were directly transferred to a pre-cooled 7ml douncer containing 3ml of ice-cold Buffer A and gently homogenized with 2 rounds, each consisting of 15 dounces over a 5-minute period. The homogenized solution was then filtered using a 37μm reversible strainer (STEMCELL 27215/27250) into a pre-cooled 15ml tube containing 3ml Buffer B. Subsequently, the mixture was centrifuged at 1000g for 30 minutes at 4°C in a swing-out rotor after inverting the tube 6 times. Careful removal of the supernatant was followed by resuspension of the pellet in 1000μl Wash Buffer, which was then filtered through a 37μm reversible strainer into a 1.5ml low-binding Axygen tube (MCT-150-L-C). Centrifugation at 500g for 5 minutes at 4°C was performed, and the wash step was repeated once more. The pellet was then resuspended in 0.5ml Buffer C and incubated on ice for 5 minutes to permeabilize nuclei. Subsequently, 0.5ml Buffer D was added and gently mixed, and the nuclei were counted. Concurrently, a final centrifugation at 500g for 5 minutes at 4°C was carried out, followed by resuspension in Buffer E at a targeted concentration of 8M nuclei/ml for loading onto the 10X Genomic chip.

**Buffer ingredient table**
| **Ingredient** | **Required Final conc** | **Unit of Required Final conc** | **Ingredient** | **Required Final conc** | **Unit of Required Final conc** |
| --- | --- | --- | --- | --- | --- |
| **Buffer A (Lysis buffer)** |  |  | **Buffer C** |  |  |
| Sucrose | 250 | mM | NaCl | 10 | mM |
| KCl | 25 | mM | MgCl<sub>2</sub> | 3 | mM |
| MgCl<sub>2</sub> | 5 | mM | Tris-HCl, pH 7.5 | 10 | mM |
| Tris-HCl, pH 7.5 | 20 | mM | NP 40 | 0.01 | % |
| Igepal CA-630 | 0.1 | % | Tween-20 | 0.01 | % |
| Spermidine | 0.5 | mM | Digitonin | 0.001 | % |
| Spermine | 0.15 | mM | Protease inhibitor | 1 | X |
| Protease inhibitor | 1 | X | DTT | 1 | mM |
| DTT | 1 | mM | BSA | 1 | % |
| Rnase Inhibitor | 0.08 | U/ul | Rnase Inhibitor | 1 | U/ul |
| **BufferB**<br>(Iodixanol) |  |  | **Buffer D** |  |  |
| KCl | 25 | mM | NaCl | 10 | mM |
| MgCl<sub>2</sub> | 5 | mM | MgCl<sub>2</sub> | 3 | mM |
| Tris-HCl, pH 7.5 | 20 | mM | Tris-HCl, pH 7.5 | 10 | mM |
| Optiprep<br>(Iodixanol) | 50 | % | Tween-20 | 0.1 | % |
| BSA+10x Protease<br>Inhibitor | 1 | % | Rnase Inhibitor | 1 | U/ul |
| DTT | 1 | mM | DTT | 1 | mM |
| Rnase Inhibitor | 0.08 | U/ul | BSA | 1 | % |
| **Wash Buffer** |  |  | **Buffer E** |  |  |
| NaCl | 10 | mM | 20X Nuclei buffer<br>(10xGenomics) | 1 | X |
| MgCl<sub>2</sub> | 3 | mM | DTT | 1 | mM |
| Tris-HCl, pH 7.5 | 10 | mM | Rnase Inhibitor | 1 | U/ul |
| Rnase Inhibitor | 1 | U/ul |  |  |  |
| DTT | 1 | mM | All the Buffers were prepared in DNase, Rnase-free H<sub>2</sub>O |  |  |
| BSA | 1 | % |  |  |  |

### Single-cell Analysis and Quality Control

Single-cell RNA sequencing (scRNA-seq) reads obtained from HOTT-derived cells, along with their corresponding primary tumor cells, were aligned to the GRCh38 human reference genome. Cell-by-gene count matrices were generated using the 10XGenomics Cell Ranger pipeline with default parameters. Subsequently, data analysis was conducted using the Seurat R package within R Studio.Cells expressing a minimum of 1000 genes and exhibiting less than 10% mitochondrial gene content were retained. Unique Molecular Identifier (UMI) counts were normalized via logarithmic transformation, employing a scaling factor of 10000. Principal Component Analysis (PCA) was executed on the scaled data, utilizing the top 2000 variable genes. The number of dimensions (dims) was determined as has been previously described (Shekhar et al., 2016). Briefly, we selected significant PCs from the larger value between the square of the standard deviation of PCA scores (Seurat.Obj@reductions$pca@stdev^2) and the square of the square root of the ratio of the number of genes to the number of cells plus one (sqrt(length(row.names(Seurat.Obj)) / length(colnames(Seurat.Obj))) + 1)^2). Cells were clustered in the PCA space utilizing Seurat’s FindNeighbors and FindClusters functions, with a resolution parameter set to 2.0. Visualization of cells was performed using Uniform Manifold Approximation and Projection (UMAP), employing the previously defined dimensions. To predict and exclude doublets, the DoubletFinder R package (McGinnis et al., 2019) was employed with default parameters.

### Copy Number Variation Analysis

To mitigate the risk of false positive or negative identification of tumor cells via FACS, copy number variation analysis was performed Infercnv with default parameters (Tickle T, 2019). Batch-matched naïve organoids were applied as reference.

### Single-cell Annotation

To perform tumor cell annotation, a projection to meta-atlas approach was employed to mitigate technical variations. The GBM meta-atlas was constructed by integrating a total of 69,547 cells from seven previously published primary GBM single-cell RNA sequencing (scRNAseq) datasets, which also included normal cells from adjacent regions proximal to the tumor, identified in the original studies (Bhaduri et al., 2020b; Couturier et al., 2020; Darmanis et al., 2017; Jacob et al., 2020; Neftel et al., 2019b; Yu et al., 2020; Yuan et al., 2018). Data analysis was performed using the Seurat R package with identical parameters as specified, except for the resolution parameter, which was set to 0.5. Subsequently, 23 clusters generated by the pipeline underwent biological annotation based on both comprehensive literature review of top-expressing genes and spatial relationships. A UMAP containing a total of 14 typical GBM cell types was generated as a reference for subsequent neoplastic cell annotation in this study.

Upon generation of Seurat query objects from HOTT datasets and reference object from GBM meta-atlas independently, FindTransferAnchors() was employed to identify a set of anchors between them, followed by MapQuery() to map the query data onto the reference in its UMAP space. Consequently, cell type annotation presented as predicted.id was generated alongside predicted.id.score to indicate the fidelity of such defined cell types to the typical GBM cell types delineated in the GBM meta-atlas. HOTT normal cell annotation was conducted in a similar manner using a recently generated meta-atlas (Nano et al., 2023) constructed from seven previously published transcriptomic profiles of the developing human cortex.

### Immunofluorescence staining

The human cortical organoids were fixed in 4% paraformaldehyde for 45 minutes at room temperature. Subsequently, the cortical organoids were rinsed with PBS and equilibrated in 30% sucrose in PBS overnight at 4C. After equilibration, the cortical organoids were embedded in OCT Embedding Matrix (Fisher Sci 14-373-65) containing OCT (Tissue-Tek, VWR) and 30% sucrose at a 1:1 ratio and frozen on a sheet of dry ice. The frozen blocks were either transferred to -80°C for long-term storage or cryosectioned as 10-16 um-thick sections onto glass slides for immunofluorescence staining. To set up the immunostaining, the sections were first rinsed with PBS for 15 minutes and then treated with a citrate-based antigen unmasking solution (10 mM sodium citrate, pH 6, Vector Labs) that was heated to 95C for 20 minutes. Following antigen unmasking, the sections were permeabilized and blocked with a blocking buffer containing 5% donkey serum, 3% bovine serum albumin, and 0.1% Triton X-100 in PBS for 30 minutes at room temperature. The subsequent primary antibody incubations were performed overnight at 4C using the following primary antibodies in the blocking buffer. Mouse -SOX2 (1:500, Santa Cruz SC-365823), mouse -Nestin (1:500, Millipore Sigma MAB5326), chicken -GFP (1:500, Aves GFP-1020), chicken -GFAP (1:500, Abcam AB4674), rabbit -Ki67 (1:500, Abcam AB16667), rabbit -MEF2C (1:500, Novus NBP1-89468), rabbit -HOPX (1:500, Proteintech 11419-1-AP). The primary antibody incubations were followed by three 10-minute PBS washes and secondary antibody incubations with DAPI in the blocking buffer for 2 hours at room temperature. The following secondary antibodies were used. DAPI (1:1000, ThermoScientific 62248), AlexaFluor 488 donkey -chicken (1:500, Jackson 703-545-155), AlexaFluor 488 donkey -goat (1:500, Invitrogen A32814), AlexaFluor 568 donkey -mouse (1:500, Invitrogen A32773), AlexaFluor 568 donkey -rat (1:500, Invitrogen A78946), AlexaFluor 568 donkey -chicken (1:500, Invitrogen A11041), AlexaFluor 647 donkey -rabbit (1:500, Invitrogen A31573). Finally, the slides were mounted with ProLongTM Gold antifade reagent (Invitrogen) and preserved for imaging at 4C.

### Image Analysis

The image acquisition parameters (light, exposure, and digital gain) were normalized based on the parameters of the images with the highest fluorescence intensities for each channel. 9 fields of fluorescent images were captured for each condition using EVOS M5000 digital-inverted benchtop microscope and individual channels were saved as TIFF files. ImageJ was used to merge the individual channels, and Imaris image analysis software (Bitplane) was used to quantify the number of double-positive cells expressing GFP and another marker of interest. These quantitative analyses were performed using the Spots and Surfaces features in Imaris. First, the GFP+ cells were determined based on a standardized shortest distance between DAPI+ Spots and GFP Surfaces. Subsequently, double-positive cells were identified based on another standardized shortest distance between GFP+ cells and either Spots or Surfaces representing a specific marker of interest.

### Tumor Cell Type Heterogeneity Preservation Analysis

Primary tumor cells were individually analyzed using the Seurat pipeline with default parameters, followed by cluster marker gene identification based on annotated tumor cell types. Subsequently, the gene score for each gene was calculated by multiplying its average log2FC by the ratio of pct.1 to pct.2, followed by pivoting to generate one gene score matrix (cluster by gene) for each primary tumor. The gene score matrices for corresponding HOTT tumor cells from each culture condition were generated using a similar approach, with the exception that cluster marker genes were identified based on Seurat-derived clusters. Gene score matrices from each culture condition within a tumor sample were merged by gene, resulting in the HOTT tumor cluster gene score matrix. The correlation between each primary tumor cell type gene score matrix and its corresponding HOTT tumor cluster gene score matrix was performed. The resulting correlation matrices of 3 primary tumor samples was then merged into a correlation matrix comprising 205 HOTT-derived clusters against primary tumor cell types, which was used to generate a heatmap using Morpheus via hierarchical clustering and visualized using a relative color scheme.

### ssGSEA Analysis

Datasets profiling HOTT cells derived from each of the three human tumors (LB4878, LB5024 and LB5028) were merged and each gene was assigned a gene score across the three samples. The Seurat clusters were separated based on culture conditions and each cluster was compared to fourteen gene lists of different cell types using ssGSEA2 (Barbie et al., 2009; Subramanian et al., 2005). The gene lists were obtained from the GBM meta-atlas. A FDR p-value threshold of less than 0.01 was applied to the clusters and their corresponding enrichment scores were plotted on a ridgeplot.

### Module Preservation Analysis

The preservation of normal developmental gene programs in primary versus transplanted tumor cells was compared using our previously identified set of gene meta-modules representative of normal human brain development (Nano et al., 2023). These 225 meta-modules were defined using iterative, hierarchical clustering of seven previously published transcriptomic profiles of the developing human cortex, and biological processes were assigned to these meta-modules through extensive literature review and term enrichment analysis (Nano et al., 2023). We then characterized the preservation of these normal developmental meta-modules in our datasets of primary and transplanted tumor cells by applying our module activity score. This versatile metric is measured for each cell by calculating the average normalized counts per million (CPM) detected for each meta-module gene. Heatmap generation and visualization of the subsequent results were conducted using Morpheus.

### Differential Gene Expression Analysis

Differential Gene Expression analysis was performed using the Seurat package Wilcoxon rank sum test with a p-value threshold of less than 0.05. Gene Ontology analysis was conducted using EnrichR.

### Cell Chat Analysis

The CellChat package (Jin et al., 2021) was employed to assess cell-to-cell communication according to its standard protocol, as outlined in https://github.com/sqjin/CellChat. Each tumor sample, combined from various culture conditions within HOTTs, underwent individual analysis using its corresponding Seurat object as input for CellChat. Samples were stratified based on both neoplastic (denoted by “N” followed by their predicted.id) and non-neoplastic (denoted by “T” followed by their predicted.id) cell types. Visualization of signaling pathway ranking was facilitated through netAnalysis_signalingRole_heatmap. Additionally, PTN receptor contribution plots, signaling pathway network circle plots, and pathway expression pattern 2D plots were generated using netAnalysis_contribution, netVisual_aggregate, and netAnalysis_signalingRole_scatter, respectively, after assigning signaling to “PTN”.

### Core versus Periphery Tumor Resection

Acquisition of glioblastoma specimens was approved under Institutional Review Board #10-000655. Patients undergoing surgery for newly diagnosed glioma were prospectively identified and completed signed informed consent. All patients had standard of care magnetic resonance imaging (MRI) (Kaufmann et al., 2020) sequences with and without gadolinium contrast (Gd-DTPA, 0.1 mmol/kg) and T2 sequences within 4 weeks of surgery. Inclusion criteria included: treatment naïve, IDH wild-type glioblastoma, contrast-enhancing bulk of tumor with surrounding non-enhancing T2 hyperintense region on magnetic resonance imaging (MRI), no peripheral involvement of eloquent areas, and medical stability for surgery. As previously described (Patel et al., 2024; Patel et al., 2020), MRI based biopsy targets were selected using intraoperative neuronavigation (BrainLab Curve, BrainLab Surgical Navigation System, Munich, Germany). Contrast-enhancing tumor specimens (Core region) were first removed during standard of care resection followed by acquisition of non-enhancing infiltrating tumor specimens (Periphery region), confirmed by pathologic review with hematoxylin-eosin stains. 2 Specimens were immediately transferred to the research laboratory for further experiments.

### RNAScope and Quantification

The in situ hybridization assay was conducted using the ACD RNAscope™ HiPlex12 Reagent Kit on FFPE primary GBM samples, followed by acquisition of RNAscope images using the ZEISS LSM 880 confocal system. The 63x confocal microscopy images were processed and visualized using the 3D view feature in Imaris. The total DAPI+ cell number was then determined using automatic spot detection. Subsequently, images with the negative control probes were taken as a reference to assess with high autofluorescence and false positivity, and PTN and PTPRZ1 positive cells were manually counted. Cells with few fluorescent speckles or dim signals were not counted as positive. Finally, the percentage of positive cells was calculated by taking the ratio of manually counted positive cells over the total cell number. The examiner remained blinded to the expected or hypothesized results.

### Gliomasphere Culture and Maintenance

Gliomaspheres were maintained in a media containing DMEM/F12 (Gibco), B27 minus Vitamin A (Gibco), Penicillin-Streptomycin (Gibco), GlutaMAX (Gibco), supplemented with heparin (5 μg/mL, Sigma), EGF (20 ng/mL, Sigma), and FGF (20 ng/mL, Sigma). The cells were cultured under 37°C and 5% CO2 and routinely tested negative for mycoplasma contamination.

### Knockdown in Organoids and Gliomaspheres with Validation

The pLKO.1 plasmid used to generate the shRNA was obtained from addgene (Plasmid #10878). The puroR cassette was excised and replaced with either mCherry or EGFP, and the hPGK promoter immediately preceding the EGFP was also replaced with a CMV promoter, both via NEBuilder HiFi DNA Assembly reaction (New England Biolabs E2621L). The scramble (5’-cctaaggttaagtcgccctcg-3’) and PTPRZ1-targeting sequence (5’-gaactcacatctgagcattgt-3’) were inserted into the backbone with the manufacturer’s instructions. To generate lentivirus, these plasmids were transfected into HEK293T cells alongside psPAX2 and pMD2.G at a mass ratio of 1 pMD2.G : 3 psPAX2 : 4 pLKO.1 using Lipofectamine 2000 (Invitrogen 11668019), and media containing the lentivirus was concentrated with Lenti-X Concentrator (Takara Bio 631232) per the manufacturer’s instructions. The concentrated lentivirus was used to infect UCLA6 hESCs at a concentration of 1:200 and 1:1000 Polybrene Infection Reagent (Millipore Sigma TR-1003-G) during split. Real-time PCR and western blotting were used to validate PTPRZ1 KD.

### Migration Assay

Gliomasphere cells were dissociated into single cells with TrypLE (ThermoFisher 12605010) and transplanted onto UCLA6 organoids at 500,000 cells/organoid via the hanging drop approach. After 36 hours, the GS-transplanted organoids were allowed to fuse with a tumor-naive, age and line-matched UCLA6 organoid in an ultra-low attachment 24-well plate, with each well harboring one fusion pair. The fusions were left undisturbed for 5 days before the first media change occurred. The fusions were maintained for 3-4 weeks before being harvested for FACS and immunostaining analyses. Part of the fusions were severed with a scalpel and dissociated into single cells with Papain in preparation for FACS analysis on the Biorad S3e Cell Sorter. The migration percentage was calculated by dividing the number of GFP+ cells on the migrated side with the total number of GFP+ cells in the fusion (n=3). The remaining fusions were fixed with 4% PFA for 45 minutes and rehydrated with 30% sucrose overnight at 4C. The organoids were then embedded in OCT and sectioned for GFP immunostaining. Images were acquired using the ZEISS LSM 880 confocal system. 15 to 36 tile images were captured using a 10x lens to incorporate the entire pair of merging organoids. To determine the tumor cell coverage over the entire fusion, eight bins of identical size were applied to each fusion, with the fusion interface in the middle. The amount of GFP+ tumor cells was quantified in each bin using Imaris.

### Western Blot

Samples were lysed with RIPA buffer (Millipore Sigma R2078) containing cOmplete™ Protease Inhibitor Cocktail (Roche 04693116001) and PhosSTOP (Roche 4906845001). Protein concentrations were determined using Pierce™ BCA Protein Assay (ThermoFisher 23225). Laemmli sample buffer (BioRad 1610747) was added to the lysate and the mixture was heated at 95C for 10 minutes. 30 ug of protein was loaded in each lane of a 4– 15% Mini-PROTEAN® TGX™ Precast Protein Gels (BioRad 4561083). After SDS-PAGE, the proteins were transferred onto a nitrocellulose membrane and blocked with 5% milk for 1 hour at room temperature. Primary antibodies were added to a solution containing 2.5% Bovine Serum Albumin and 0.02% sodium azide in TBST; anti-PTPRZ1 (1:500, BD Biosciences 610179), anti-β-Actin (1:10000, Sigma-Aldrich A1978), anti-SRC (1:10000, Abcam ab109381), anti-phospho-SRC (1:1000, Cell Signaling Tech 6943T). The membranes were incubated overnight at 4C. Secondary antibodies targeting mouse (1:10000, Fisher Scientific NC9994806) or rabbit (1:10000, Fisher Scientific NC9734651) in 5% milk was added to the membranes for 1 hour at room temperature. The membranes were then developed with Immobilon Forte Western HRP substrate (Millipore Sigma WBLUF).

### Quantitative Real-time PCR

RNA extraction was performed with RNeasy Plus Mini Kit (Qiagen 74136) according to the manufacturer’s manual and the resulting RNA was used to synthesize cDNA using SuperScript™ IV VILO™ Master Mix (Invitrogen 11756050). Real-time PCR targeting PTPRZ1 (5’-ACTCTGAGAAGCAGAGGAG-3’ and 5’-CTGTTGTCTGTAGTATCCATTAG-3’) or GAPDH (5’-TCAAGGCTGAGAACGGGAAG-3’ and 5’-CGCCCCACTTGATTTTGGAG-3’) was performed with Power SYBR™ Green PCR Master Mix (Applied Biosystems 4367659) in three technical replicates.

**SFig1:**
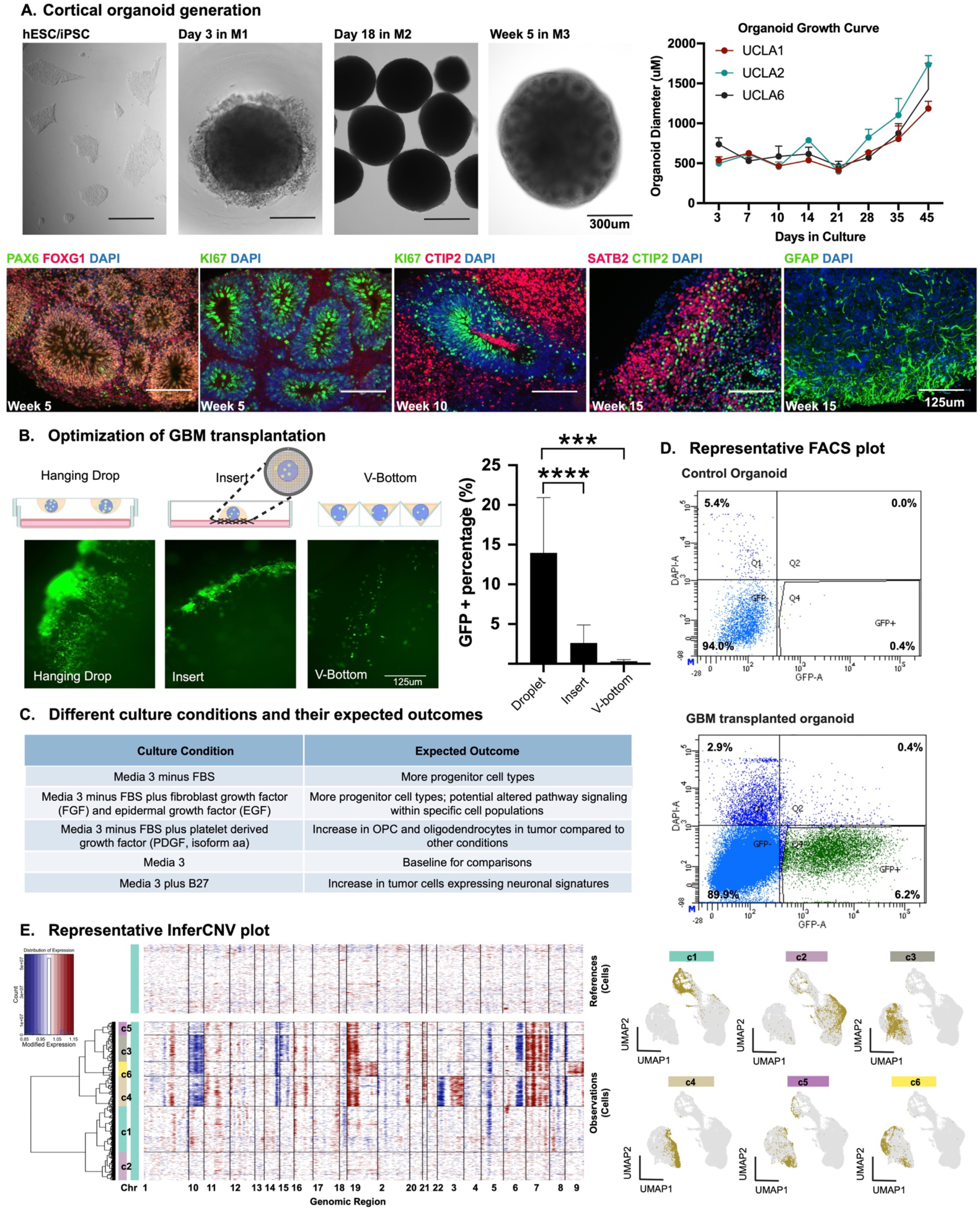
Experimental Setup for <u>H</u>uman <u>O</u>rganoid <u>T</u>umor <u>T</u>ransplantation (HOTT) model (Associated with Fig 1) **A)** Cortical organoids were derived from multiple lines of hESC (human embryonic stem cells). Top panel: Representative images of cortical organoids cultivated in various differentiation media, together with a growth curve presented to track their size growth. To generate cortical organoids, human embryonic stem cells were dissociated into single cell suspensions and aggregated into ultra-low attachment V-bottom plates in the presence of ROCK, WNT and TGFb inhibitors (Media 1, M1). These organoids were introduced into Media 2 (M2) and Media 3 (M3) on Day 18 and Day 35, respectively, to further shape them towards a cortical fate. Bottom panel: IF verification illustrating the stepwise progression towards cortical fate. At Week 5, neural progenitor characteristics were evident, characterized by the presence of rosette structures and positive staining for PAX6 and Ki67. Subsequent neural differentiation was marked by the expression of the deep-layer neuronal marker CTIP2 at Week 10, followed by the appearance of the upper-layer neuronal marker SATB2 and the typical astrocytic marker GFAP at Week 15. **B)** Three different transplantation methods of HOTT were compared using the same quantity of tumor cells. After three weeks of culturing in M3 media, all transplanted organoids were sectioned and stained for GFP (Green Fluorescent Protein), and the percentage of GFP-positive area relative to the total organoid area in the field was calculated to assess transplantation efficiency. **C)** Five different culture conditions were employed to investigate the response of GBM to various environmental perturbations, with expected outcomes outlined alongside each condition. **D)** Following 3-4 weeks of culture in different media, FACS (Fluorescence-Activated Cell Sorting) sorting was performed to enrich GFP-labeled GBM cells. A gate was established using GFP-negative (GFP-) naive organoids (top panel), and only samples with over 1000 GFP-positive (GFP+) cells proceeded to single-cell transcriptome analysis. **E)** InferCNV was used to identify neoplastic cells from non-neoplastic cells, as shown by the representative InferCNV plots and feature plots to highlight neoplastic cell clusters corresponding to InferCNV clusters.

**SFig2:**
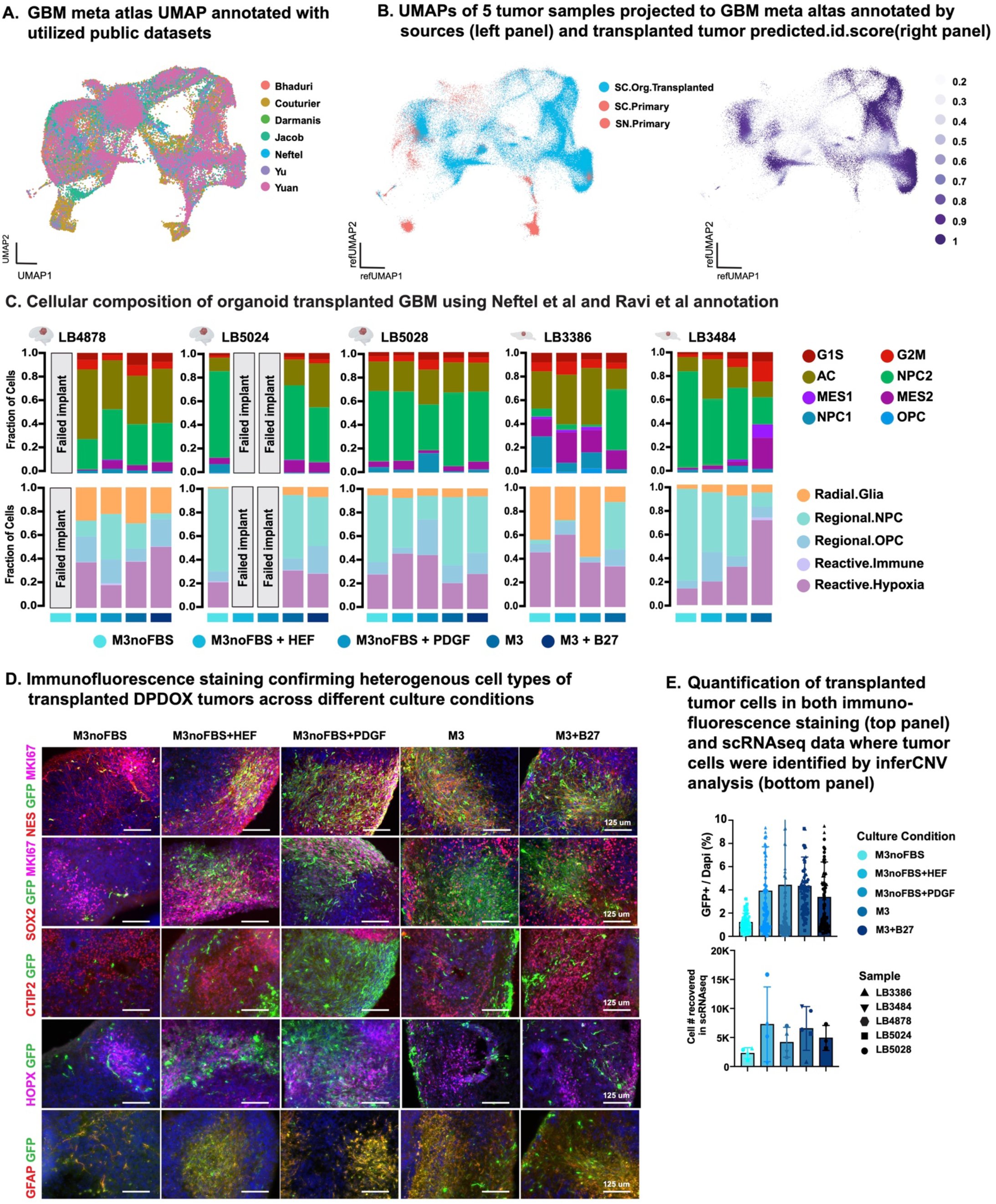
Characterization of Tumor Heterogeneity in HOTT System (Associated with Figures 1 and 2) **A)** A compendium of 7 GBM published datasets was used to generate a GBM meta-atlas. These datasets were downloaded from publicly available data depositions and filtered to maintain only tumor cells as indicated by copy number variation analysis. Integration was performed using RPCA analysis (Seurat v4) and clusters were annotated for cell type. The UMAP shown here depicts the integrated analysis yielded co-mingling of all 7 datasets included. **B)** The projection of 5 HOTT GBM samples alongside their corresponding primary tumors onto the GBM meta-atlas facilitates the reduction of batch effects and technical disparities between single-cell and single-nucleus RNAseq methodologies. In projected UMAP space annotated by source (left panel), the preservation of the majority of the tumor complexity was shown with less maintenance of certain cell types such as microglia and oligodendrocytes, which are only present in primary tumor samples in red. While in projected UMAP space annotated by predicted ID score (right panel), high molecular similarity was shown between HOTT cell types and their corresponding GBM meta-atlas cell types. **C)** Cellular composition of 5 HOTT GBM samples utilizing Neftel et al and Ravi et al annotations, complementing Fig 2A, not only exhibited similar trends to those observed with the predicted ID annotation but also showed a robust correlation among the major cell types annotated using three different annotation systems. **D)** Additional immunofluorescence validation of the most populous cell types within transplanted DPDOX tumors across various culture conditions, with their quantification displayed in Fig 2B and 2C. **E)** Additional quantification was conducted to assess tumor cell numbers in 5 HOTT samples across 5 different culture conditions. The number of GFP+ cells in immunofluorescence (IF) assays matched the tumor cell count obtained via single-cell RNAseq following identification through inferCNV features. Consistently, FBS-containing culture conditions exhibited higher tumor cell counts, whereas base media without FBS yielded the fewest tumor cells. Moreover, supplementation with growth factors (HEF and PDGF) increased tumor cell numbers.

**SFig 3.**
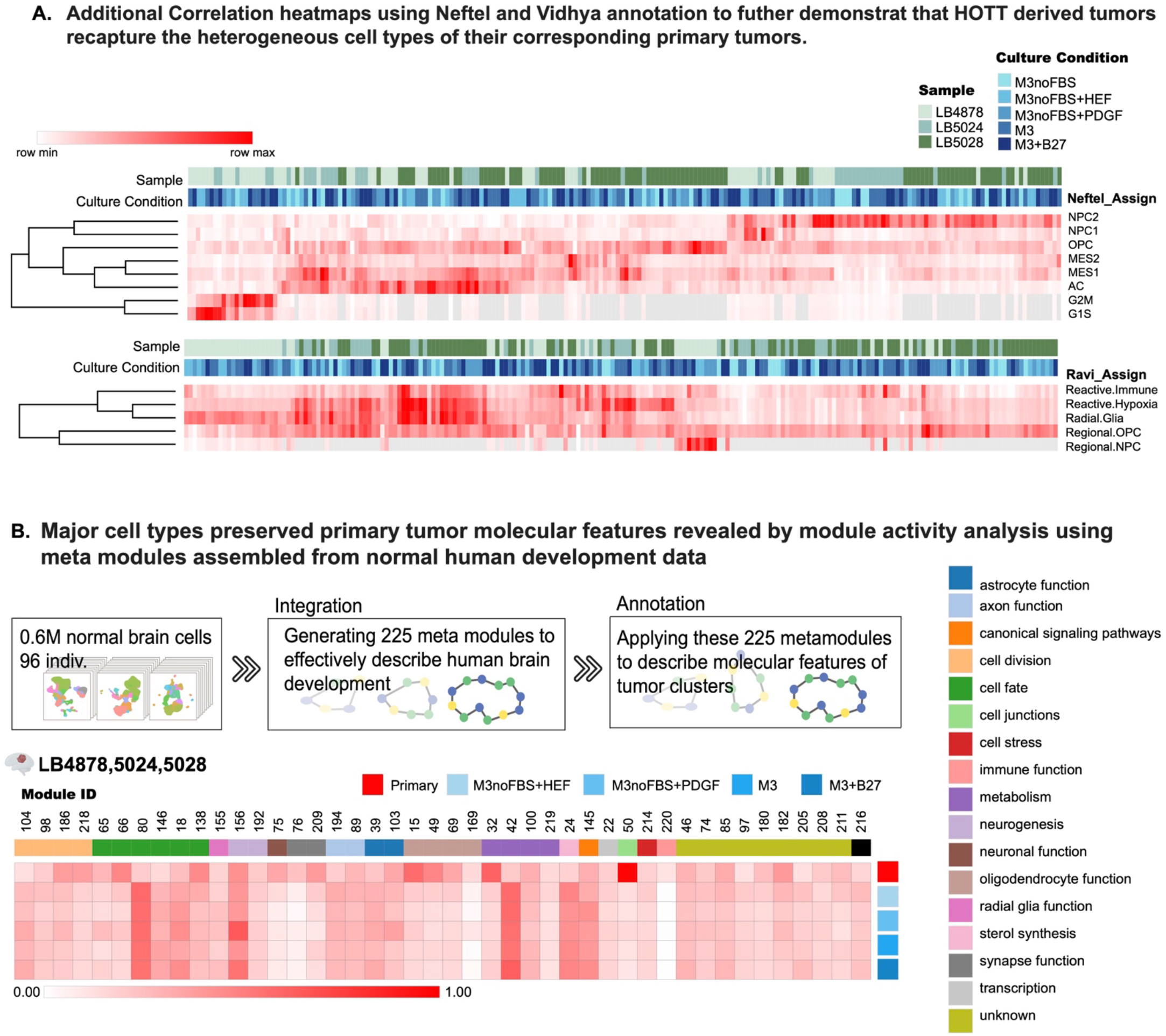
HOTT derived tumor cells recapitulate the molecular features of their corresponding primary tumors. (Associated with Figure 3) **A)** Additional heatmaps demonstrate the correlation of HOTT derived tumor cells to their corresponding primary tumors using Neftel et al and Ravi et al annotations further demonstrates the recapitulation of tumor cell type heterogeneity. **B)** Characterization of tumor cells based on the activity of gene modules representing normal human brain development. Gene modules were identified by applying a novel strategy of iterative, hierarchical clustering to a meta-atlas of seven previously published transcriptomic profiles of the developing human cortex (Nano et al., 2023) (top). This pipeline produced 225 modules, which were assigned biological annotations through both extensive literature review of module genes and term enrichment analysis of gene ontology sets, WikiPathway and KEGG pathway databases, and ChEA, ENCODE and TRRUST transcriptional regulatory collections. A module activity score based on average module gene expression was calculated for each tumor cell. Heat map (bottom) shows the average module activity in primary cells or in transplanted tumor cells across three samples grown in the indicated culture conditions. When compared to their corresponding primary tumors, most HOTT tumors maintained comparable module activity, except for immune function, cell junction, and synapse functions.

**SFig 4.**
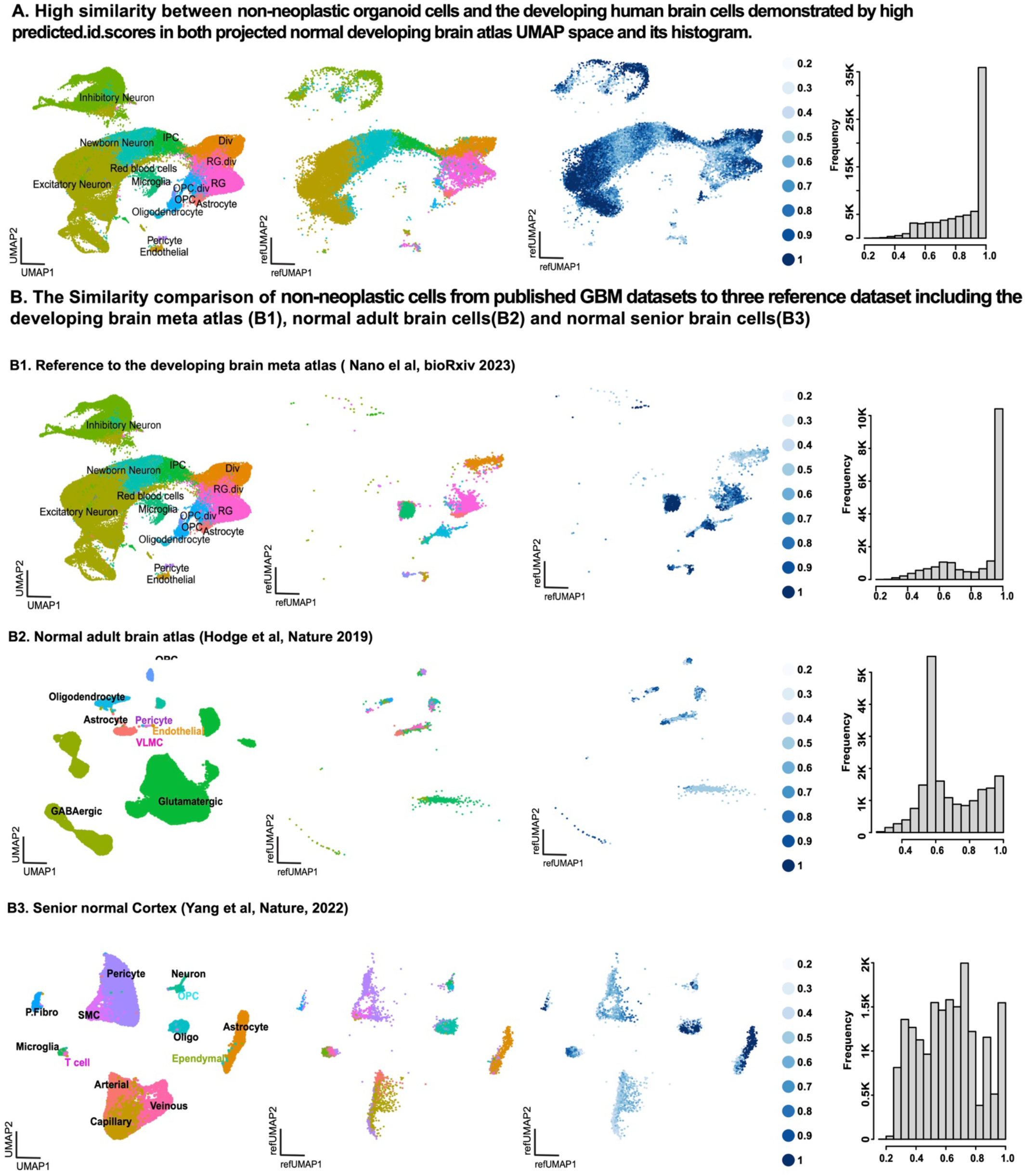
Non-neoplastic cells derived from both HOTT organoids and published GBM dataset exhibited the greatest similarity to corresponding cell types found in the developing human brain. (Associated with Figure 4) To investigate the interaction of tumor cells and their surrounding InferCNV-identified non-neoplastic cells. Non-neoplastic cell types were annotated using three different reference databases: normal developing brain meta-atlas (A and B1), adult brains (B2), and normal aging brains (B3). Corresponding UMAPs displayed in the left column. **A)** As expected, we observed the highest predicted scores of HOTT non-neoplastic organoid cells when projected to the developing brain meta-atlas, as demonstrated by their UMAP in the reference UMAP space annotated by cell types (predicted.id), the similarity (predicted.id.score) and histogram of mean predicted scores, respectively. **B)** When annotating the non-neoplastic cells from published GBM datasets, the highest predicted scores were also observed when projected to the developing brain meta-atlas (B1), but not to normal adult brain cells (B2) and normal aging brain cells(B3). The results suggest the applicability of using the HOTT system to study the tumor microenvironment.

**SFig 5.**
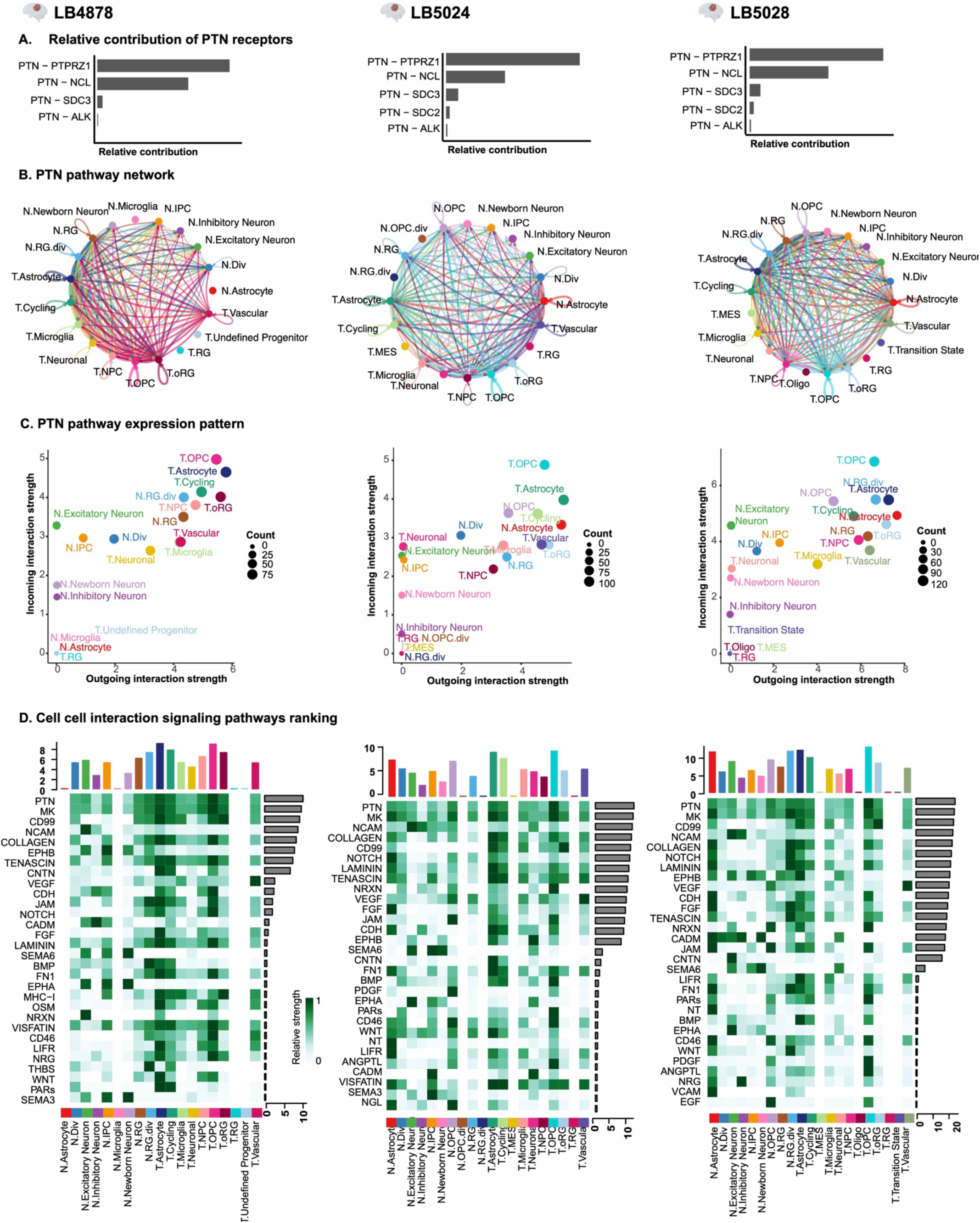
Additional CellChat analysis on three HOTT samples across culture conditions yielded consistent results, highlighting the role of PTN-PTPRZ1 signaling in mediating interactions between the tumor and its microenvironment.

**SFig 6.**
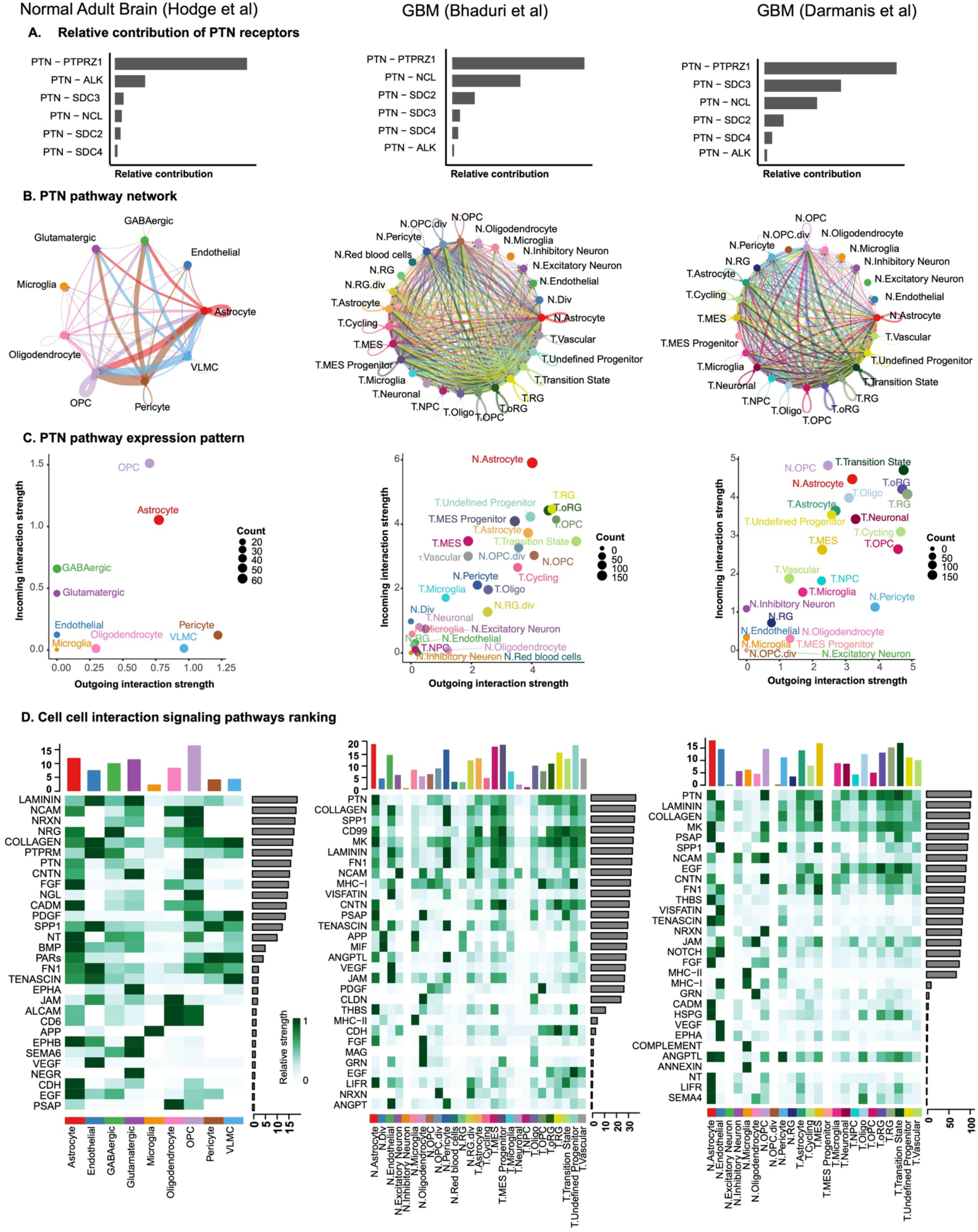
Additional CellChat analysis on published adult normal brain and GBM datasets with non-neoplastic cells confirmed PTN/PTPRZ1 signaling as the top pathway mediating the interactions in tumor microenvironment but not in normal brain environment.

**SFig 7.**
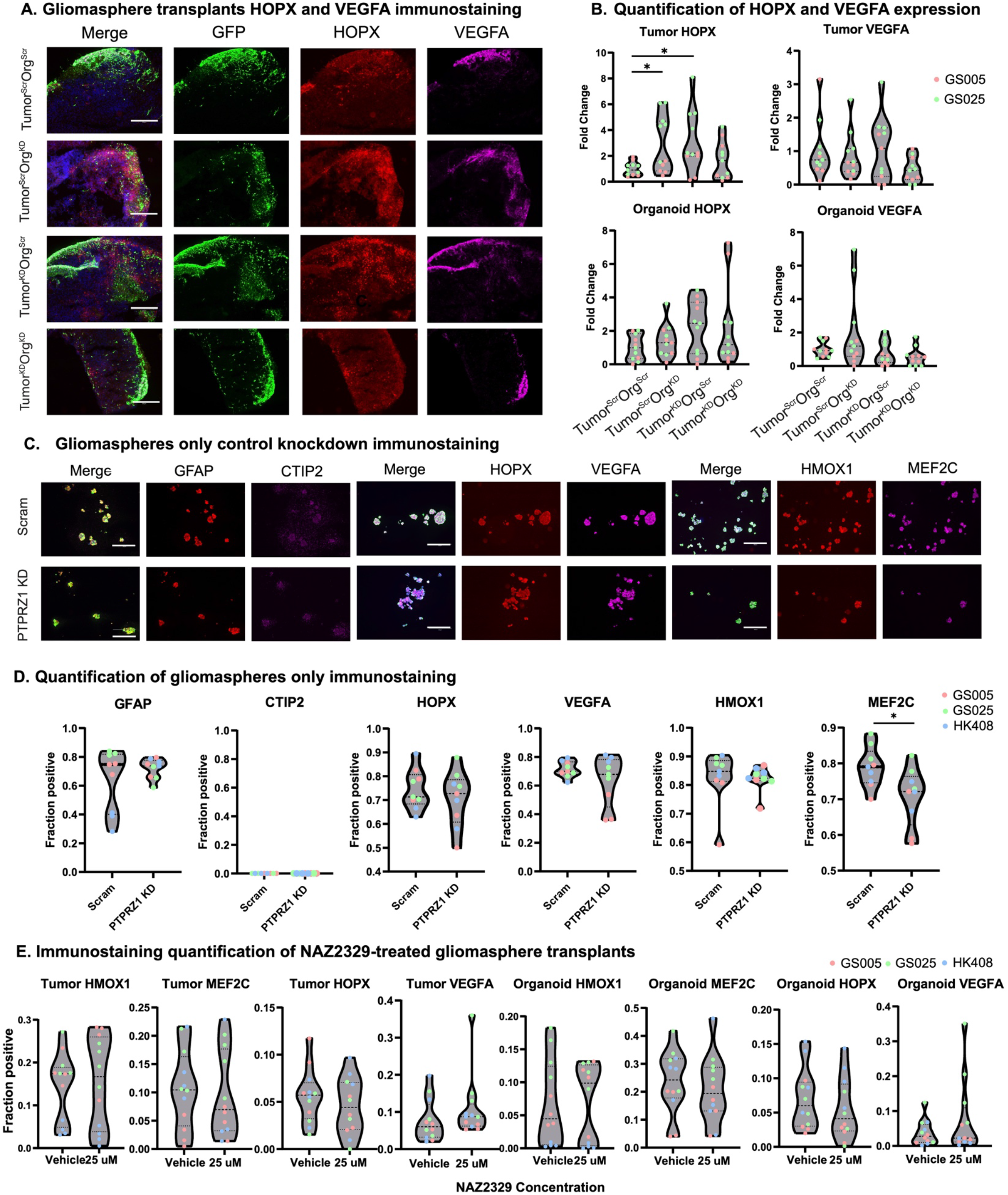
The tumor microenvironment is essential to PTPRZ1-mediated cell type changes. **A)** Effects of tumor or environmental PTPRZ1 KD on cell fate were assessed with immunostaining of radial glial marker HOPX and mesenchymal marker VEGFA. Tumor KD of PTPRZ1 significantly increases the expression of tumor HOPX and organoid HOPX, suggesting that intrinsic and environmental PTPRZ1 both play a role in suppressing radial glia fate in the tumor. **B)** Violin plots depicting the quantified tumor and organoid expression of HOPX and VEGFA. Fold change of each experimental condition was calculated based on the average of all the data points in Tumor^Scr^Org^Scr^. Tumor and organoid expression were based on the ratio of GFP+/Signal of interest+ over total GFP+ cells and GFP-/Signal of interest+ over total GFP-cells, respectively. P-value determined by two-tailed Mann-Whitney test, *p ≤ 0.05. Experiments were conducted in two patient derived gliomasphere lines (GS005 and GS025) with a minimum of 3 but up to 6 replicates per line. **C)** Immunostaining of astrocyte marker GFAP, neuronal markers CTIP2 and MEF2C, radial glia marker HOPX, and mesenchymal markers HMOX1 and VEGFA of gliomaspheres only under scramble or PTPRZ1 KD conditions (n=3). Modulation in cell fate has largely disappeared upon PTPRZ1 KD, highlighting that the presence of a tumor microenvironment is essential for the observed cell type changes. **D)** Violin plots depicting the quantified immunostaining in D. P-value determined by two-tailed Mann-Whitney test, *p ≤ 0.05. Experiments were conducted in three patient derived gliomasphere lines (GS005, GS025 and HK408) with 3 replicates per line. **E)** Violin plots depicting the quantified expression of HMOX1, MEF2C, HOPX, and VEGFA of GS transplanted organoids under vehicle and 25 μM NAZ2329 treatment. No significant cell type changes were observed, indicating that cell fate specification was not driven by PTPRZ1 catalytic activity. Experiments were conducted in three GS lines (GS005, GS025 and HK408) with 4 replicates per line. P-value determined by two-tailed Mann-Whitney test.

## References Cited

Akter, F., Simon, B., de Boer, N.L., Redjal, N., Wakimoto, H., and Shah, K. (2021). Pre-clinical tumor models of primary brain tumors: Challenges and opportunities. Biochim Biophys Acta Rev Cancer 1875, 188458.

Bagley, S.J., Kothari, S., Rahman, R., Lee, E.Q., Dunn, G.P., Galanis, E., Chang, S.M., Nabors, L.B., Ahluwalia, M.S., Stupp, R., et al. (2022). Glioblastoma Clinical Trials: Current Landscape and Opportunities for Improvement. Clin Cancer Res 28, 594–602.

Bakken, T.E., Jorstad, N.L., Hu, Q., Lake, B.B., Tian, W., Kalmbach, B.E., Crow, M., Hodge, R.D., Krienen, F.M., Sorensen, S.A., et al. (2021). Comparative cellular analysis of motor cortex in human, marmoset and mouse. Nature 598, 111–119.

Barbie, D.A., Tamayo, P., Boehm, J.S., Kim, S.Y., Moody, S.E., Dunn, I.F., Schinzel, A.C., Sandy, P., Meylan, E., Scholl, C., et al. (2009). Systematic RNA interference reveals that oncogenic KRAS-driven cancers require TBK1. Nature 462, 108–112.

Bhaduri, A., Andrews, M.G., Mancia Leon, W., Jung, D., Shin, D., Allen, D., Jung, D., Schmunk, G., Haeussler, M., Salma, J., et al. (2020a). Cell stress in cortical organoids impairs molecular subtype specification. Nature 578, 142–148.

Bhaduri, A., Di Lullo, E., Jung, D., Muller, S., Crouch, E.E., Espinosa, C.S., Ozawa, T., Alvarado, B., Spatazza, J., Cadwell, C.R., et al. (2020b). Outer Radial Glia-like Cancer Stem Cells Contribute to Heterogeneity of Glioblastoma. Cell Stem Cell 26, 48–63 e46.

Cadwell, C.R., Bhaduri, A., Mostajo-Radji, M.A., Keefe, M.G., and Nowakowski, T.J. (2019). Development and Arealization of the Cerebral Cortex. Neuron 103, 980–1004.

Couturier, C.P., Ayyadhury, S., Le, P.U., Nadaf, J., Monlong, J., Riva, G., Allache, R., Baig, S., Yan, X., Bourgey, M., et al. (2020). Single-cell RNA-seq reveals that glioblastoma recapitulates a normal neurodevelopmental hierarchy. Nature communications 11, 3406.

Darmanis, S., Sloan, S.A., Croote, D., Mignardi, M., Chernikova, S., Samghababi, P., Zhang, Y., Neff, N., Kowarsky, M., Caneda, C., et al. (2017). Single-Cell RNA-Seq Analysis of Infiltrating Neoplastic Cells at the Migrating Front of Human Glioblastoma. Cell Rep 21, 1399–1410.

de Visser, K.E., and Joyce, J.A. (2023). The evolving tumor microenvironment: From cancer initiation to metastatic outgrowth. Cancer Cell 41, 374–403.

Fujikawa, A., Nagahira, A., Sugawara, H., Ishii, K., Imajo, S., Matsumoto, M., Kuboyama, K., Suzuki, R., Tanga, N., Noda, M., et al. (2016). Small-molecule inhibition of PTPRZ reduces tumor growth in a rat model of glioblastoma. Sci Rep 6, 20473.

Fujikawa, A., Sugawara, H., Tanaka, T., Matsumoto, M., Kuboyama, K., Suzuki, R., Tanga, N., Ogata, A., Masumura, M., and Noda, M. (2017). Targeting PTPRZ inhibits stem cell-like properties and tumorigenicity in glioblastoma cells. Sci Rep 7, 5609.

Gupta, K., and Burns, T.C. (2018). Radiation-Induced Alterations in the Recurrent Glioblastoma Microenvironment: Therapeutic Implications. Front Oncol 8, 503.

Hansen, D.V., Lui, J.H., Parker, P.R., and Kriegstein, A.R. (2010). Neurogenic radial glia in the outer subventricular zone of human neocortex. Nature 464, 554–561.

Hasselbach, L.A., Irtenkauf, S.M., Lemke, N.W., Nelson, K.K., Berezovsky, A.D., Carlton, E.T., Transou, A.D., Mikkelsen, T., and deCarvalho, A.C. (2014). Optimization of high grade glioma cell culture from surgical specimens for use in clinically relevant animal models and 3D immunochemistry. J Vis Exp, e51088.

Hodge, R.D., Bakken, T.E., Miller, J.A., Smith, K.A., Barkan, E.R., Graybuck, L.T., Close, J.L., Long, B., Johansen, N., Penn, O., et al. (2019). Conserved cell types with divergent features in human versus mouse cortex. Nature 573, 61–68.

Jacob, F., Salinas, R.D., Zhang, D.Y., Nguyen, P.T.T., Schnoll, J.G., Wong, S.Z.H., Thokala, R., Sheikh, S., Saxena, D., Prokop, S., et al. (2020). A Patient-Derived Glioblastoma Organoid Model and Biobank Recapitulates Inter- and Intra-tumoral Heterogeneity. Cell 180, 188–204 e122.

Janiszewska, M., Suva, M.L., Riggi, N., Houtkooper, R.H., Auwerx, J., Clement-Schatlo, V., Radovanovic, I., Rheinbay, E., Provero, P., and Stamenkovic, I. (2012). Imp2 controls oxidative phosphorylation and is crucial for preserving glioblastoma cancer stem cells. Genes Dev 26, 1926–1944.

Jin, S., Guerrero-Juarez, C.F., Zhang, L., Chang, I., Ramos, R., Kuan, C.H., Myung, P., Plikus, M.V., and Nie, Q. (2021). Inference and analysis of cell-cell communication using CellChat. Nat Commun 12, 1088.

Jorstad, N.L., Close, J., Johansen, N., Yanny, A.M., Barkan, E.R., Travaglini, K.J., Bertagnolli, D., Campos, J., Casper, T., Crichton, K., et al. (2023). Transcriptomic cytoarchitecture reveals principles of human neocortex organization. Science 382, eadf6812.

Jourdon, A., Wu, F., Mariani, J., Capauto, D., Norton, S., Tomasini, L., Amiri, A., Suvakov, M., Schreiner, J.D., Jang, Y., et al. (2023). Modeling idiopathic autism in forebrain organoids reveals an imbalance of excitatory cortical neuron subtypes during early neurogenesis. Nat Neurosci 26, 1505–1515.

Kadoshima, T., Sakaguchi, H., Nakano, T., Soen, M., Ando, S., Eiraku, M., and Sasai, Y. (2013). Self-organization of axial polarity, inside-out layer pattern, and species-specific progenitor dynamics in human ES cell-derived neocortex. Proc Natl Acad Sci U S A 110, 20284–20289.

Kaufmann, T.J., Smits, M., Boxerman, J., Huang, R., Barboriak, D.P., Weller, M., Chung, C., Tsien, C., Brown, P.D., Shankar, L., et al. (2020). Consensus recommendations for a standardized brain tumor imaging protocol for clinical trials in brain metastases. Neuro Oncol 22, 757–772.

Krawczyk, M.C., Haney, J.R., Pan, L., Caneda, C., Khankan, R.R., Reyes, S.D., Chang, J.W., Morselli, M., Vinters, H.V., Wang, A.C., et al. (2022). Human Astrocytes Exhibit Tumor Microenvironment-, Age-, and Sex-Related Transcriptomic Signatures. J Neurosci 42, 1587–1603.

Krishna, S., Choudhury, A., Keough, M.B., Seo, K., Ni, L., Kakaizada, S., Lee, A., Aabedi, A., Popova, G., Lipkin, B., et al. (2023). Glioblastoma remodelling of human neural circuits decreases survival. Nature 617, 599–607.

Lathia, J.D., Mack, S.C., Mulkearns-Hubert, E.E., Valentim, C.L., and Rich, J.N. (2015). Cancer stem cells in glioblastoma. Genes Dev 29, 1203–1217.

LeBlanc, V.G., Trinh, D.L., Aslanpour, S., Hughes, M., Livingstone, D., Jin, D., Ahn, B.Y., Blough, M.D., Cairncross, J.G., Chan, J.A., et al. (2022). Single-cell landscapes of primary glioblastomas and matched explants and cell lines show variable retention of inter- and intratumor heterogeneity. Cancer Cell 40, 379–392 e379.

Lee, J., Kotliarova, S., Kotliarov, Y., Li, A., Su, Q., Donin, N.M., Pastorino, S., Purow, B.W., Christopher, N., Zhang, W., et al. (2006). Tumor stem cells derived from glioblastomas cultured in bFGF and EGF more closely mirror the phenotype and genotype of primary tumors than do serum-cultured cell lines. Cancer Cell 9, 391–403.

Linkous, A., Balamatsias, D., Snuderl, M., Edwards, L., Miyaguchi, K., Milner, T., Reich, B., Cohen-Gould, L., Storaska, A., Nakayama, Y., et al. (2019). Modeling Patient-Derived Glioblastoma with Cerebral Organoids. Cell Rep 26, 3203–3211 e3205.

Lui, J.H., Hansen, D.V., and Kriegstein, A.R. (2011). Development and evolution of the human neocortex. Cell 146, 18–36.

Mancusi, R., and Monje, M. (2023). The neuroscience of cancer. Nature 618, 467–479.

Mathys, H., Peng, Z., Boix, C.A., Victor, M.B., Leary, N., Babu, S., Abdelhady, G., Jiang, X., Ng, A.P., Ghafari, K., et al. (2023). Single-cell atlas reveals correlates of high cognitive function, dementia, and resilience to Alzheimer’s disease pathology. Cell 186, 4365–4385 e4327.

McGinnis, C.S., Murrow, L.M., and Gartner, Z.J. (2019). DoubletFinder: Doublet Detection in Single-Cell RNA Sequencing Data Using Artificial Nearest Neighbors. Cell Syst 8, 329–337 e324.

Meng, K., Rodriguez-Pena, A., Dimitrov, T., Chen, W., Yamin, M., Noda, M., and Deuel, T.F. (2000). Pleiotrophin signals increased tyrosine phosphorylation of beta beta-catenin through inactivation of the intrinsic catalytic activity of the receptor-type protein tyrosine phosphatase beta/zeta. Proc Natl Acad Sci U S A 97, 2603–2608.

Mikelis, C., Sfaelou, E., Koutsioumpa, M., Kieffer, N., and Papadimitriou, E. (2009). Integrin alpha(v)beta(3) is a pleiotrophin receptor required for pleiotrophin-induced endothelial cell migration through receptor protein tyrosine phosphatase beta/zeta. FASEB J 23, 1459–1469.

Muthukrishnan, S.D., Kawaguchi, R., Nair, P., Prasad, R., Qin, Y., Johnson, M., Wang, Q., VanderVeer-Harris, N., Pham, A., Alvarado, A.G., et al. (2022). P300 promotes tumor recurrence by regulating radiation-induced conversion of glioma stem cells to vascular-like cells. Nat Commun 13, 6202.

Nano, P.R., Fazzari, E., Azizad, D., Nguyen, C.V., Wang, S., Kan, R.L., Wick, B., Haeussler, M., and Bhaduri, A. (2023). A Meta-Atlas of the Developing Human Cortex Identifies Modules Driving Cell Subtype Specification. bioRxiv.

Neftel, C., Laffy, J., Filbin, M.G., Hara, T., Shore, M.E., Rahme, G.J., Richman, A.R., Silverbush, D., Shaw, M.L., Hebert, C.M., et al. (2019a). An Integrative Model of Cellular States, Plasticity, and Genetics for Glioblastoma. Cell.

Neftel, C., Laffy, J., Filbin, M.G., Hara, T., Shore, M.E., Rahme, G.J., Richman, A.R., Silverbush, D., Shaw, M.L., Hebert, C.M., et al. (2019b). An Integrative Model of Cellular States, Plasticity, and Genetics for Glioblastoma. Cell 178, 835–849 e821.

Omuro, A., and DeAngelis, L.M. (2013). Glioblastoma and other malignant gliomas: a clinical review. JAMA 310, 1842–1850.

Osswald, M., Jung, E., Sahm, F., Solecki, G., Venkataramani, V., Blaes, J., Weil, S., Horstmann, H., Wiestler, B., Syed, M., et al. (2015). Brain tumour cells interconnect to a functional and resistant network. Nature 528, 93–98.

Ostrem, B., Di Lullo, E., and Kriegstein, A. (2017). oRGs and mitotic somal translocation - a role in development and disease. Curr Opin Neurobiol 42, 61–67.

Ostrem, B.E., Lui, J.H., Gertz, C.C., and Kriegstein, A.R. (2014). Control of outer radial glial stem cell mitosis in the human brain. Cell Rep 8, 656–664.

Pajonk, F., Vlashi, E., and McBride, W.H. (2010). Radiation resistance of cancer stem cells: the 4 R’s of radiobiology revisited. Stem Cells 28, 639–648.

Pastrana, E., Silva-Vargas, V., and Doetsch, F. (2011). Eyes wide open: a critical review of sphere-formation as an assay for stem cells. Cell Stem Cell 8, 486–498.

Patel, K.S., Yao, J., Cho, N.S., Sanvito, F., Tessema, K., Alvarado, A., Dudley, L., Rodriguez, F., Everson, R., Cloughesy, T.F., et al. (2024). pH-Weighted amine chemical exchange saturation transfer echo planar imaging visualizes infiltrating glioblastoma cells. Neuro Oncol 26, 115–126.

Patel, K.S., Yao, J., Raymond, C., Yong, W., Everson, R., Liau, L.M., Nathanson, D., Kornblum, H., Wang, C., Oughourlian, T., et al. (2020). Decorin expression is associated with predictive diffusion MR phenotypes of anti-VEGF efficacy in glioblastoma. Sci Rep 10, 14819.

Pollen, A.A., Bhaduri, A., Andrews, M.G., Nowakowski, T.J., Meyerson, O.S., Mostajo-Radji, M.A., Di Lullo, E., Alvarado, B., Bedolli, M., Dougherty, M.L., et al. (2019). Establishing Cerebral Organoids as Models of Human-Specific Brain Evolution. Cell 176, 743–756 e717.

Pollen, A.A., Nowakowski, T.J., Chen, J., Retallack, H., Sandoval-Espinosa, C., Nicholas, C.R., Shuga, J., Liu, S.J., Oldham, M.C., Diaz, A., et al. (2015). Molecular identity of human outer radial glia during cortical development. Cell 163, 55–67.

Pombo Antunes, A.R., Scheyltjens, I., Duerinck, J., Neyns, B., Movahedi, K., and Van Ginderachter, J.A. (2020). Understanding the glioblastoma immune microenvironment as basis for the development of new immunotherapeutic strategies. Elife 9.

Qin, E.Y., Cooper, D.D., Abbott, K.L., Lennon, J., Nagaraja, S., Mackay, A., Jones, C., Vogel, H., Jackson, P.K., and Monje, M. (2017). Neural Precursor-Derived Pleiotrophin Mediates Subventricular Zone Invasion by Glioma. Cell 170, 845–859 e819.

Ravi, V.M., Will, P., Kueckelhaus, J., Sun, N., Joseph, K., Salie, H., Vollmer, L., Kuliesiute, U., von Ehr, J., Benotmane, J.K., et al. (2022). Spatially resolved multi-omics deciphers bidirectional tumor-host interdependence in glioblastoma. Cancer Cell 40, 639–655 e613.

Shekhar, K., Lapan, S.W., Whitney, I.E., Tran, N.M., Macosko, E.Z., Kowalczyk, M., Adiconis, X., Levin, J.Z., Nemesh, J., Goldman, M., et al. (2016). Comprehensive Classification of Retinal Bipolar Neurons by Single-Cell Transcriptomics. Cell 166, 1308–1323 e1330.

Shi, Y., Ping, Y.F., Zhou, W., He, Z.C., Chen, C., Bian, B.S., Zhang, L., Chen, L., Lan, X., Zhang, X.C., et al. (2017). Tumour-associated macrophages secrete pleiotrophin to promote PTPRZ1 signalling in glioblastoma stem cells for tumour growth. Nat Commun 8, 15080.

Subramanian, A., Tamayo, P., Mootha, V.K., Mukherjee, S., Ebert, B.L., Gillette, M.A., Paulovich, A., Pomeroy, S.L., Golub, T.R., Lander, E.S., et al. (2005). Gene set enrichment analysis: a knowledge-based approach for interpreting genome-wide expression profiles. Proc Natl Acad Sci U S A 102, 15545–15550.

Tickle T, T.I., Georgescu C, Brown M, Haas B (2019). inferCNV of the Trinity CTAT Project.

Tirosh, I., Izar, B., Prakadan, S.M., Wadsworth, M.H., 2nd, Treacy, D., Trombetta, J.J., Rotem, A., Rodman, C., Lian, C., Murphy, G., et al. (2016). Dissecting the multicellular ecosystem of metastatic melanoma by single-cell RNA-seq. Science 352, 189–196.

Velasco, S., Kedaigle, A.J., Simmons, S.K., Nash, A., Rocha, M., Quadrato, G., Paulsen, B., Nguyen, L., Adiconis, X., Regev, A., et al. (2019). Individual brain organoids reproducibly form cell diversity of the human cerebral cortex. Nature 570, 523–527.

Venkataramani, V., Yang, Y., Schubert, M.C., Reyhan, E., Tetzlaff, S.K., Wissmann, N., Botz, M., Soyka, S.J., Beretta, C.A., Pramatarov, R.L., et al. (2022). Glioblastoma hijacks neuronal mechanisms for brain invasion. Cell 185, 2899–2917 e2831.

Venkatesh, H.S., Morishita, W., Geraghty, A.C., Silverbush, D., Gillespie, S.M., Arzt, M., Tam, L.T., Espenel, C., Ponnuswami, A., Ni, L., et al. (2019). Electrical and synaptic integration of glioma into neural circuits. Nature 573, 539–545.

Venteicher, A.S., Tirosh, I., Hebert, C., Yizhak, K., Neftel, C., Filbin, M.G., Hovestadt, V., Escalante, L.E., Shaw, M.L., Rodman, C., et al. (2017). Decoupling genetics, lineages, and microenvironment in IDH-mutant gliomas by single-cell RNA-seq. Science 355.

Wakimoto, H., Kesari, S., Farrell, C.J., Curry, W.T., Jr., Zaupa, C., Aghi, M., Kuroda, T., Stemmer-Rachamimov, A., Shah, K., Liu, T.C., et al. (2009). Human glioblastoma-derived cancer stem cells: establishment of invasive glioma models and treatment with oncolytic herpes simplex virus vectors. Cancer Res 69, 3472–3481.

Wang, L., Jung, J., Babikir, H., Shamardani, K., Jain, S., Feng, X., Gupta, N., Rosi, S., Chang, S., Raleigh, D., et al. (2022). A single-cell atlas of glioblastoma evolution under therapy reveals cell-intrinsic and cell-extrinsic therapeutic targets. Nat Cancer 3, 1534–1552.

Wang, R., Sharma, R., Shen, X., Laughney, A.M., Funato, K., Clark, P.J., Shpokayte, M., Morgenstern, P., Navare, M., Xu, Y., et al. (2020). Adult Human Glioblastomas Harbor Radial Glia-like Cells. Stem Cell Reports 14, 338–350.

Wen, P.Y., Weller, M., Lee, E.Q., Alexander, B.M., Barnholtz-Sloan, J.S., Barthel, F.P., Batchelor, T.T., Bindra, R.S., Chang, S.M., Chiocca, E.A., et al. (2020). Glioblastoma in adults: a Society for Neuro-Oncology (SNO) and European Society of Neuro-Oncology (EANO) consensus review on current management and future directions. Neuro Oncol 22, 1073–1113.

Winkler, F., Venkatesh, H.S., Amit, M., Batchelor, T., Demir, I.E., Deneen, B., Gutmann, D.H., Hervey-Jumper, S., Kuner, T., Mabbott, D., et al. (2023). Cancer neuroscience: State of the field, emerging directions. Cell 186, 1689–1707.

Xia, Z., Ouyang, D., Li, Q., Li, M., Zou, Q., Li, L., Yi, W., and Zhou, E. (2019). The Expression, Functions, Interactions and Prognostic Values of PTPRZ1: A Review and Bioinformatic Analysis. J Cancer 10, 1663–1674.

Yang, A.C., Vest, R.T., Kern, F., Lee, D.P., Agam, M., Maat, C.A., Losada, P.M., Chen, M.B., Schaum, N., Khoury, N., et al. (2022). A human brain vascular atlas reveals diverse mediators of Alzheimer’s risk. Nature 603, 885–892.

Yu, K., Hu, Y., Wu, F., Guo, Q., Qian, Z., Hu, W., Chen, J., Wang, K., Fan, X., Wu, X., et al. (2020). Surveying brain tumor heterogeneity by single-cell RNA-sequencing of multi-sector biopsies. Natl Sci Rev 7, 1306–1318.

Yuan, J., Levitin, H.M., Frattini, V., Bush, E.C., Boyett, D.M., Samanamud, J., Ceccarelli, M., Dovas, A., Zanazzi, G., Canoll, P., et al. (2018). Single-cell transcriptome analysis of lineage diversity in high-grade glioma. Genome Med 10, 57.

Zhu, Q., Zhao, X., Zheng, K., Li, H., Huang, H., Zhang, Z., Mastracci, T., Wegner, M., Chen, Y., Sussel, L., et al. (2014). Genetic evidence that Nkx2.2 and Pdgfra are major determinants of the timing of oligodendrocyte differentiation in the developing CNS. Development 141, 548–555.

